# A rotational velocity estimate constructed through visuomotor competition updates the fly’s neural compass

**DOI:** 10.1101/2023.09.25.559373

**Authors:** Brad K. Hulse, Angel Stanoev, Daniel B. Turner-Evans, Johannes D. Seelig, Vivek Jayaraman

## Abstract

Navigating animals continuously integrate velocity signals to update internal representations of their directional heading and spatial location in the environment. How neural circuits combine sensory and motor information to construct these velocity estimates and how these self-motion signals, in turn, update internal representations that support navigational computations are not well understood. Recent work in *Drosophila* has identified a neural circuit that performs angular path integration to compute the fly’s head direction, but the nature of the velocity signal is unknown. Here we identify a pair of neurons necessary for angular path integration that encode the fly’s rotational velocity with high accuracy using both visual optic flow and motor information. This estimate of rotational velocity does not rely on a moment-to-moment integration of sensory and motor information. Rather, when visual and motor signals are congruent, these neurons prioritize motor information over visual information, and when the two signals are in conflict, reciprocal inhibition selects either the motor or visual signal. Together, our results suggest that flies update their head direction representation by constructing an estimate of rotational velocity that relies primarily on motor information and only incorporates optic flow signals in specific sensorimotor contexts, such as when the motor signal is absent.

## Introduction

Flexible navigation requires combining diverse streams of sensory and motor information to estimate one’s movements through the world. This task is complicated by the fact that these sensorimotor streams are often noisy, unreliable, or context-dependent, leading to cue conflicts that require resolution (Noel and Angelaki, 2022). For example, when an animal turns right, the world appears to move leftwards across the eye, producing a reafferent optic flow signal that can be used to assess the magnitude of the intended turn (Gibson, 1950). Yet, when objects pass leftward in front of a stationary animal, a similar optic flow signal could be generated, even though the animal isn’t moving. It is widely recognized that experiences like these can lead to perceived self-motion (Mach, 1875; Tschermak, 1931), such as when sitting on a stationary train and seeing an adjacent train begin to move. In general, the reliability and utility of different self-motion cues depends on sensorimotor context. Optic flow would be largely absent when walking in darkness or riding a stationary bike, for instance, yet motor efference and proprioceptive signals would still be available. Given these complexities, how animals combine diverse sensorimotor streams to accurately compute self-motion estimates is incompletely understood (reviewed in (Brooks and Cullen, 2019; Chiappe, 2023; Cullen, 2019; Currier and Nagel, 2020; Huston and Jayaraman, 2011; Noel and Angelaki, 2022)).

Understanding how animals construct accurate self-motion estimates is also key to dissecting navigation computations (Cullen and Taube, 2017; Laurens and Angelaki, 2018; Savelli and Knierim, 2019). Flexible navigation requires animals to form internal representations of their spatial relationship to the environment (Fisher, 2022; Grieves and Jeffery, 2017), and integration of self-motion cues is central to updating these representations in both vertebrates (Bassett and Taube, 2001; Blair et al., 1998; Graham et al., 2023; Kropff et al., 2015; Sharp et al., 2001; Stackman and Taube, 1998) and invertebrates (Currier et al., 2020; Green et al., 2017; Hulse et al., 2021; Lu et al., 2022; Lyu et al., 2022; Matheson et al., 2022; Shiozaki et al., 2020; Stone et al., 2017; Turner-Evans et al., 2017; Weir and Dickinson, 2015). One particularly prominent internal representation supports our sense of direction and is composed of neurons known as head direction (HD) cells that encode the orientation of the head in a world-centered reference frame (reviewed in (Hulse and Jayaraman, 2019; Taube, 2007)). Originally discovered in rodents (Rank, 1984; Taube et al., 1990a), HD cells have since been reported in a diverse array of animals, including bats, birds, fish, and insects, highlighting their evolutionary importance (Ben-Yishay et al., 2021; Finkelstein et al., 2015; Heinze and Homberg, 2007; Petrucco et al., 2022; Seelig and Jayaraman, 2015; Varga and Ritzmann, 2016; Vinepinsky et al., 2020). Conceptually, when HD cells are arranged on a ring according to their preferred directional tunings, their population activity is organized as a traveling bump whose angular position encodes the animal’s HD, much like the needle on a compass (Skaggs et al., 1995). Considerable theoretical work has established that a class of networks known as ring attractors can elegantly account for these dynamics (Amari, 1977; Ben-Yishai et al., 1995; Hansel and Sompolinsky, 1998; Noorman et al., 2022; Redish et al., 1996; Zhang, 1996). Indeed, recent experimental studies in *Drosophila* have discovered a ring attractor network that encodes the fly’s HD as a bump of activity that travels around a toroidal structure known as the ellipsoid body (EB), a striking example of network structure matching function (**Figure 1A**) (Green et al., 2017; Kim et al., 2017b; Seelig and Jayaraman, 2015; Turner-Evans et al., 2017; Turner-Evans et al., 2021). To update the position of their HD bump, animals integrate rotational velocity information across time, but the nature of the velocity signal is not well understood (Cullen and Taube, 2017; Laurens and Angelaki, 2018). Understanding which sensorimotor cues are used to construct the rotational velocity signal, whether certain cues are prioritized over others, and how conflicts are resolved will shed light on how sensorimotor computations like these are tailored to the navigation computations they support.

**Figure 1:**
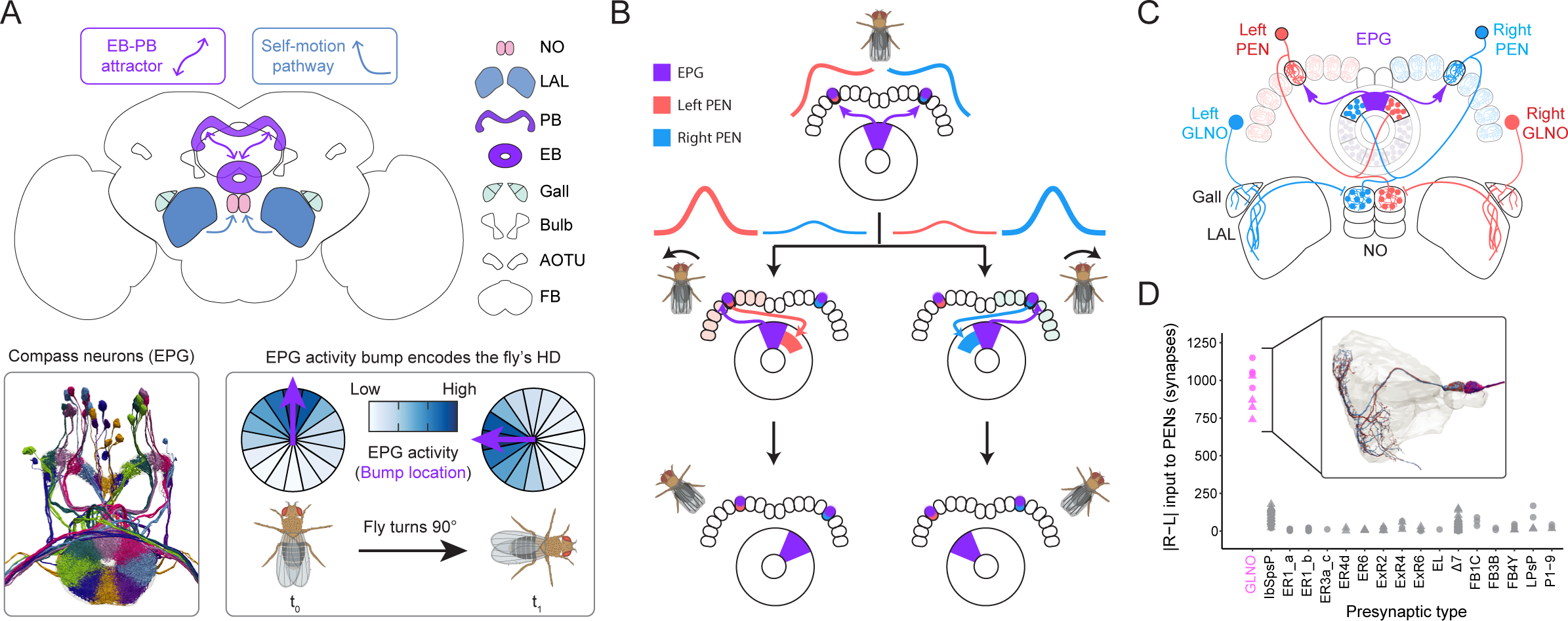
A neural circuit for angular path integration in the Drosophila central complex. **A)** Top: schematic of the fly brain with the central complex and associated neuropil outlined along with a pathway that conveys self-motion information from the LAL to the NO (blue arrows). Neuropil acronyms: noduli (NO), lateral accessory lobe (LAL), protocerebral bridge (PB), ellipsoid body (EB), anterior optic tubercle (AOTU), fan-shaped body (FB). Bottom left: morphological rendering of the fly’s compass neurons showing how individual neurons (marked by distinct colors) divide the EB into ‘wedges’ that are tuned to distinct head directions (electron-micros- copy-based reconstruction by Janelia FlyEM: (Hulse et al., 2021; Scheffer et al., 2020)). Bottom right: schematic showing that when the fly turns clockwise by 90 degrees, its head direction (HD) bump, carried by the EPG population, rotates 90 degrees counterclockwise. **B)** Schematic showing how PEN neurons (likely PEN_a: (Green et al., 2017)) update the position of the EPG bump, adapted from (Turner-Evans et al., 2017). The EPG neurons convey their bump from the EB to the left and right PB, forming two bumps that get inherited by the left (red bump) and right PEN (blue bump) populations. The PEN populations have their bump amplitudes differentially modulated by a rotational velocity signal and project back to the EB with either a clockwise or counterclockwise offset, allowing them to update the EB bump position in either direction. **C)** Schematic showing that GLNO neurons contact either the left or right PEN populations in the left and right NO, providing a potential mecha- nism for them to differentially control the amplitude of two PEN bumps. **D)** Plot showing the degree to which individual neurons upstream of the PEN neurons differentially target the left and right populations separately, a necessary requirement for generating PEN-driven updates of the EPG bump. For every individual neuron presynaptic to PEN neurons (PEN_a neurons marked by circles, PEN_b neurons by triangles), we computed the absolute difference in the number of synapses onto the right and left PEN populations. Note that GLNO neurons are the only highly lateralized input to PEN neurons. Other neuron types show a small degree of lateralization, such as the IbSpsP and Δ7 neurons, but these populations are known to target many other central complex neuron types that are not involved in angular integration.

Recent work leveraging the powerful circuit tools available in *Drosophila* has begun to provide a circuit- and cell-type level understanding of sensorimotor-driven navigation computations, setting the stage for a deeper understanding of how self-motion estimates are constructed and ultimately used to update internal representations (Currier et al., 2020; Green et al., 2017; Hulse et al., 2021; Lu et al., 2022; Lyu et al., 2022; Matheson et al., 2022; Shiozaki et al., 2020; Stone et al., 2017; Turner-Evans et al., 2017). These navigation computations are housed in a highly conserved brain region known as the central complex whose intricately woven circuits implement a variety of vector-based navigation computations (Fisher, 2022; Honkanen et al., 2019; Pfeiffer and Homberg, 2014; Turner-Evans and Jayaraman, 2016; Wilson, 2023), beginning with the computation of HD in the EB (**Figure 1A** (Hulse and Jayaraman, 2019; Seelig and Jayaraman, 2015)). Flies update their HD representation by continuously integrating rotational velocity information and by relying on directional sensory cues like visual landmarks when present (Fisher et al., 2019; Fisher et al., 2022; Haberkern et al., 2022; Kim, 2021; Kim et al., 2019; Okubo et al., 2020; Seelig and Jayaraman, 2013; Sun et al., 2017). While the source of rotational velocity information has not been established, the circuit mechanisms supporting angular velocity integration have been worked out in considerable detail and involve two main populations: EPG neurons and PEN neurons. (**Figure 1B**; (Green et al., 2017; Turner-Evans et al., 2017; Turner-Evans et al., 2020)). EPG neurons, which function as HD cells, forward their activity bump from the EB to left and right halves of a handlebar-shaped structure known as the protocerebral bridge (PB; **Figure 1B**). This generates two bumps: one in the left bridge and one in the right bridge. There, the bumps are inherited by a second population of neurons known as PEN neurons. PEN neurons from the left and right bridge project back to the EB, but with a ∼45-degree phase shift in either the clockwise or counterclockwise direction, respectively (**Figure 1B**). Importantly, the left and right PEN populations are conjunctively tuned to the fly’s HD and rotational velocity (Green et al., 2017; Turner-Evans et al., 2017). This rotational velocity signal acts to differentially control the amplitude of the left and right PEN bumps, generating appropriate compass updates when the fly turns. PEN neurons have two subtypes, PEN_a and PEN_b, each with distinct rotational tuning (Green et al., 2017) and circuit connectivity (Hulse et al., 2021; Turner-Evans et al., 2020).

Here, we address two key open questions: how do flies combine sensory and motor information to estimate their rotational velocity across different sensorimotor contexts, and how does this rotational velocity information interact with PEN neurons to generate accurate bump updates? Using whole-brain connectomics, causal circuit perturbations, and two-photon calcium imaging from flies walking in virtual reality, we identify a bilateral pair of neurons—known as GLNO neurons—as the sole source of the rotational velocity information to PEN neurons. GLNO neurons are necessary for angular path integration and their activity encodes the fly’s rotational velocity with high accuracy using both motor and visual information. The GLNO neurons on the right side of the brain are active both during right turns and in response to optic flow stimuli consistent with those generated by right turns, and the left GLNO neurons are active during left turns and in response to optic flow stimuli matching those generated by left turns. Under most circumstances, when visual and motor signals are largely congruent, GLNO neurons rely on motor information at the expense of optic flow, effectively bypassing visuomotor integration by suppressing reafferent visual feedback. Yet, when motor signals and reafferent visual feedback convey opposing velocity estimates, lateral inhibition acting upstream of GLNO neurons generates flip-flop dynamics that can resolve the visuomotor conflict by relying on one cue at the expense of the other. Given that the presence and content of visual and motor cues depends on context, our results suggest flies update their HD representation by constructing a rotational velocity estimate primarily using motor signals, but can transiently switch to relying on visual self-motion signals.

## Results

### GLNO neurons are the primary source of rotational velocity input to PEN neurons

Recent indirect evidence implicates GLNO neurons as one potential source of rotational velocity information that could update the fly’s HD representation. In addition to innervating the EB and PB, PEN neurons also innervate a paired midline neuropil known as the noduli (NO), with PEN neurons from the left bridge innervating the right NO and PEN neurons from the right bridge innervating the left NO (**Figure 1C**) (Lin et al., 2013; Wolff et al., 2015; Wolff and Rubin, 2018).

Building on these anatomical insights, a recent connectomic analysis identified a pathway that could convey self-motion information from a prominent sensorimotor center known as the lateral accessory lobe (LAL) to the NO (**Figure 1A** (Hulse et al., 2021; Namiki and Kanzaki, 2016a, b; Scheffer et al., 2020). This pathway is composed of parallel input channels formed by distinct types of so-called “LNO” neurons, each of which is thought to convey self-motion information to central complex neurons implementing distinct vector computations. Consistent with this, recent physiological recordings in bees (Stone et al., 2017) and flies (Currier et al., 2020; Lu et al., 2022; Lyu et al., 2022) have identified LNO types that convey translational self-motion information to central complex circuits that compute the fly’s direction of travel. Given that GLNO neurons are the sole LNO type targeting the PEN neurons, these results suggest that GLNO neurons may convey rotational velocity information to the HD network. In agreement with this, GLNO neurons have strong direct connections to both PEN_a and PEN_b neurons (Hulse et al., 2021), and functional connectivity experiments suggest that GLNO neurons inhibit PEN neurons (Franconville et al., 2018).

Importantly, despite these indications, it remains possible that other neurons may convey rotational velocity information to PEN neurons as well. To effectively drive bump updates, any rotational velocity signal would have to differentially control the amplitude of the left and right PEN bumps, requiring lateralized inputs that preferentially target the left and right PEN populations separately. Building on a recent connectomic analysis (Hulse et al., 2021), we searched for lateralized inputs across all upstream inputs to PEN_a and PEN_b neurons. This analysis revealed that GLNO neurons are the only highly lateralized input to PEN neuron (**Figure 1D**), suggesting that they are the only potential source of rotational velocity information for the fly’s compass network.

### GLNO neurons are required for angular path integration

If GLNO neurons are the main source of rotational velocity information for the fly’s HD network, then impairing their function should impair the network’s ability to accurately integrate rotational velocity. To test this, we reduced synaptic transmission from GLNO neurons and measured the impact on EPG bump dynamics by performing two-photon calcium imaging as flies walked on a spherical treadmill in darkness (**Figure 2A-B**) or in closed-loop with a prominent visual landmark (**Figure 2 S1**) (see Methods). GLNO synaptic transmission was reduced through expression of shibire^ts^, a temperature-sensitive mutant of *Drosophila*’s dynamin ortholog, which decreases vesicle endocytosis and therefore reduces synaptic release at elevated temperatures (Kitamoto, 2001). Flies lacking shibire^ts^ served as controls for the effect of elevated temperature on fly behavior (**Figure 2 S1 C-D**), as in previous studies (Turner-Evans et al., 2017). At room temperature, the EPG bump tracked the fly’s HD in both shibire^ts^ and control flies, consistent with previous results (Seelig and Jayaraman, 2015). In contrast, at elevated temperatures, EPG bump dynamics in shibire^ts^ flies became completely uncoupled from the fly’s HD (**Figure 2C**, bottom panel), while control flies continued to display normal bump updates (**Figure 2C**, top panel). This core result held across three GLNO driver lines and in flies walking in darkness (**Figure 2D**) or in closed loop with a prominent visual landmark (**Figure 2 S1**). Altering synaptic transmission from GLNO neurons using shibire^ts^ did not prevent the bump from moving altogether, nor did it alter the bump’s shape parameters, such as its amplitude (**Figure 2 S1 B**). Instead, it prevented the bump from accurately tracking the fly’s HD, even when a prominent visual landmark was present, consistent with the notation that forming and maintaining an accurate visual map requires accurate path integration (Kim et al., 2019).

**Figure 2:**
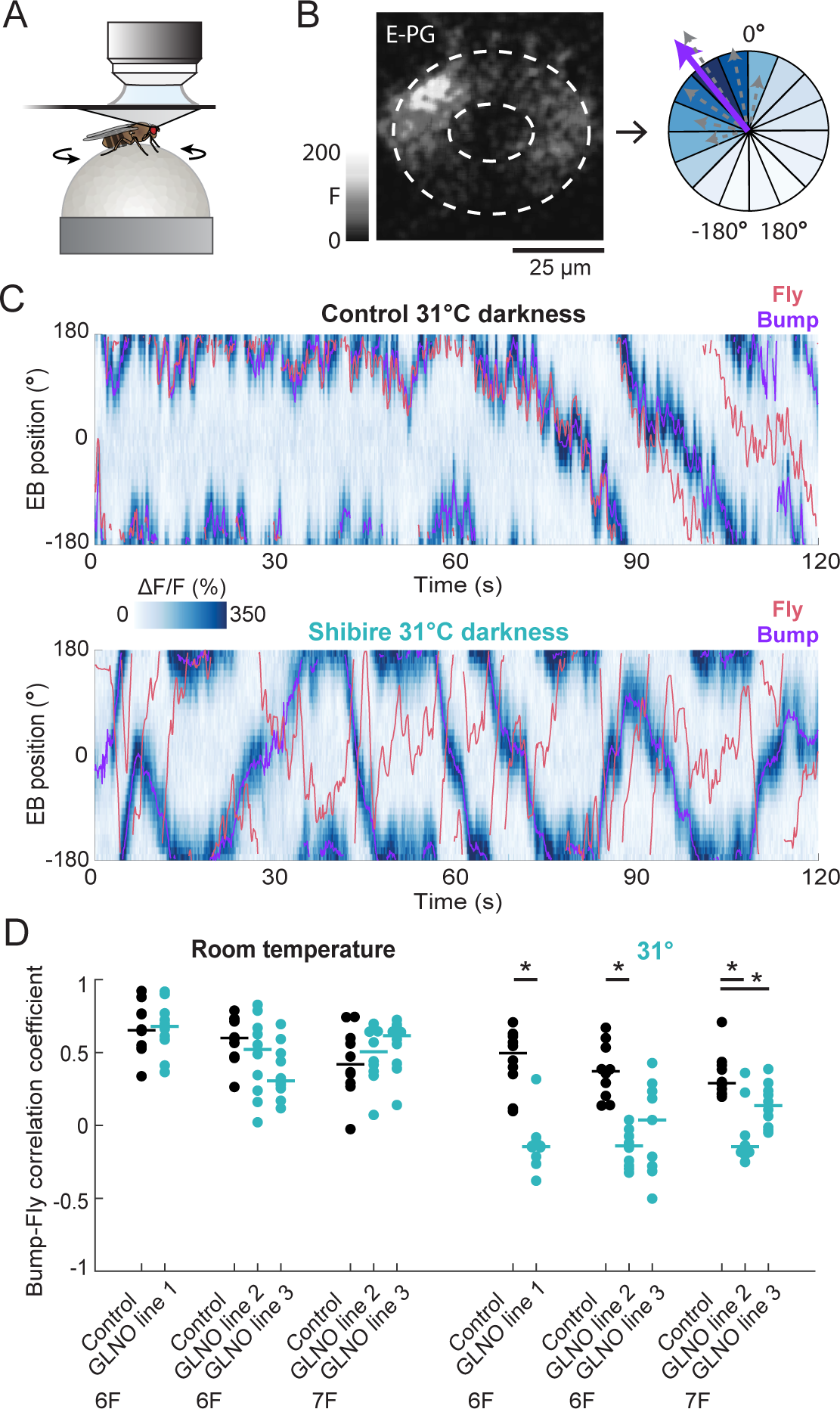
GLNO neurons are required for accurate angular integration by the fly compass. **A)** Illustration of our experimental setup, where head-fixed flies walk on an air-cushioned, spherical treadmill during two-photon calcium imaging of neural activity. A visual display (not shown) surrounds the fly (240° in azimuth and ∼55° in elevation), allowing for either closed-loop or open-loop presentation of visual stimuli. **B)** Left: snapshot of raw two-photon GCaMP7f fluorescence reflecting EPG calcium activity. Notice the single bump of activity. Right: to compute the bump’s location, the EB is divided into wedge-shaped ROIs, ΔF/F is computed for each ROI (across time), and a population vector average (PVA) is calculated (purple arrow) across ROIs (gray arrows). **C)** Top: example trial showing EPG bump dynamics from a control fly walking in darkness at 31° C. Notice that the fly’s EPG bump (purple line) tracks the fly’s HD (pink line), but with the gradual accumulation of error over time, a well-known feature of angular path integration in darkness. Bottom: same as above, but for a fly expressing shibire^ts^ in GLNO neurons. Notice that impairing synaptic release from GLNO neurons causes movements of the bump to decouple from the fly’s true HD. **D)** Quantification showing the correlation coefficient between the fly’s unwrapped HD and its unwrapped bump position for control flies (black circles) and flies expressing shibire^ts^ in GLNO neurons (green circles) using three different driver lines (GLNO Line 1-3; see Methods), both at room temperature (left half of plot) and at 31° (right half of plot), when shibire^ts^ reduced vesicle release from the GLNO neurons. Each circle is data from one fly, with the population median marked by the horizonal line. Black bars with asterisks indicate significant pairwise differences between group medians as assessed with a Wilcoxon rank sum test with multiple comparison correction using the false discovery rate (see methods). Notice that the bump-fly correlation is only reduced in flies expressing shibire^ts^ walking at 3 1° C.

### GLNO neurons encode the fly’s rotational velocity using motor information

Next, we performed two-photon calcium imaging from both the left and right GLNO neurons as flies walked in darkness to determine whether their activity encodes the fly’s rotational velocity (**Figure 3A-B**). Indeed, the right GLNO neurons were strongly active during right turns and the left GLNO neurons were strongly active during left turns, generating a “flip-flop” activity pattern (Namiki and Kanzaki, 2016a, b; Olberg, 1983) where either the left or the right GLNO neurons were highly active, though rarely at the same time (**Figure 3C**, top panel). GLNO neurons are relatively inactive during turns in their non-preferred direction, suggesting that the fly’s rotational velocity is encoded by the difference in their activity, consistent with notion that bump updates are driven by differential activation of the left and right PEN populations (Green et al., 2017; Turner-Evans et al., 2017). When we directly compared the difference in GLNO activity (right minus left) to the fly’s rotational velocity, we found the two signals are highly correlated (**Figure 3C**, bottom panel), with tuning curves that are well fit by a sigmoid (**Figure 3D**). Across flies, the difference in GLNO activity was highly correlated with the fly’s rotational velocity (**Figure 3E**), with an approximate zero-time lag (**Figure 3F**, left panel), but not with its forward velocity (**Figure 3F**, right panel). The fly-to-fly variability in tuning curve slope (**Figure 3E**) is consistent with previously reported fly-to-fly variability in the gain of angular integration (Seelig and Jayaraman, 2015) and in PEN tuning curve slopes (Turner-Evans et al., 2017).

**Figure 3:**
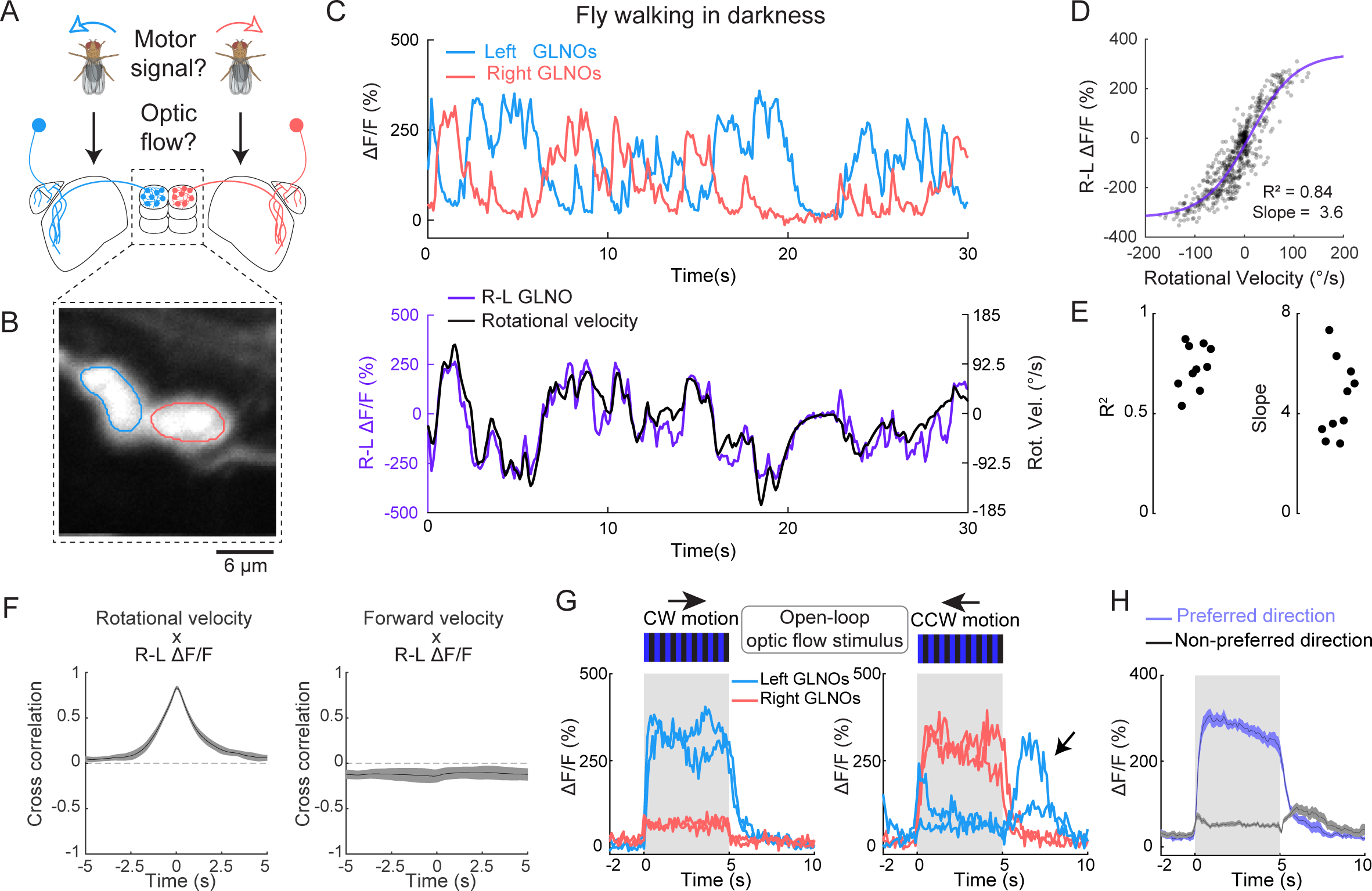
**GLNO neurons encode the fly’s rotational velocity using both a motor signal and an optic flow signal**. **A)** By recording GLNO activity from flies walking in darkness (panels C-E), we assessed whether the left GLNO neurons are active during left turns and if the right GLNO neurons are active during right turns, as suggested by a model of PEN-driven bump updates (Figure 1C). Similarly, by recording from immobilized flies viewing visual motion (panels G-H), we assessed whether GLNO neurons are also driven by optic flow stimuli. **B)** Raw two-photon GCaMP7f fluorescence of GLNO calcium activity averaged across all frames from a representative fly, with ROI outlines shown for the left (blue) and right (red) NO processes. **C)** Top: example trial showing calcium activity for the left (blue) and right (red) GLNO neurons. Notice the flip-flop activity pattern, during which either the left or the right GLNO neurons are active, but rarely at the same time. Bottom: the difference in GLNO activity (right minus left, shown in purple) is plotting alongside the fly’s rotational velocity (shown in black). As in previous studies (Turner-Evans et al., 2017), we convolved the fly’s raw rotational velocity with a kernel modeling GCaMP dynamics to compare the two signals on a similar timescale. **D)** Scatter plot showing the difference in GLNO activity (y-axis) as a function of the fly’s rotational velocity (x-axis) using the data from (C). The purple line shows a sigmoidal fit to the GLNO tuning curve. **E)** Left: beeswarm plot showing the distribution of R² values across flies for sigmoidal tuning curve fits, as in (D). Right: Same but for the maximum slope of the tuning curve. **F)** Fly-averaged (N=10) cross correlations (+/- s.e.m.) between the flies’ rotational (left panel) or forward velocity (right panel) and the R-L GLNO activity. **G)** Example GLNO data from one fly presented with a 45° square wave grating moving in either the clockwise (CW; left panel) or counterclock- wise (CW; right panel) direction at 45°/s. Notice that the left GLNO neurons, which are active during left turns, are also driven by visual motion that would arise from a left turn (CW motion), and that the right GLNO neurons, which are active during right turns, are also driven by visual motion that would arise from a right turn (CCW motion). Black arrow in the right panel marks a spontaneous activation of the left GLNO neurons. . **H)** Fly-averaged (N=10 flies) GLNO responses (+/- s.e.m.) to motion in the preferred (purple) and non-preferred (black) directions. Left GLNO neurons prefer CW motion while right GLNOs prefer CCW motion, as shown in panel G.

Together, these results support a model where GLNO neurons drive bump updates by inhibiting the PEN population tuned to contralateral turns (**Figure 1D**). That is, right GLNO neurons, which are active during right turns, inhibit PEN neurons that project to the left side of the PB, to decrease their bump amplitude during right turns. Similarly, left GLNO neurons, which are active during left turns, inhibit PEN neurons whose bump is smallest during left turns. Thus, GLNO neurons use inhibition to differentially control the amplitude of the left and right PEN bumps. Importantly, because visual cues were absent while flies walked in darkness, our results demonstrate that GLNO neurons can extract rotational velocity using motor information alone, either from proprioceptive cues or internally-generated efference copies (Chiappe, 2023; Sperry, 1950; von Holst and Mittelstaedt, 1950) which could arise from descending neuron collaterals or their upstream inputs (Rayshubskiy et al., 2020; Ros et al., 2023; Schnell et al., 2017). As noted in previous studies (Chiappe, 2023; Fenk et al., 2021; Fujiwara et al., 2022; Fujiwara et al., 2017; Kim et al., 2017a; Kim et al., 2015), untangling contributions from motor efference copies and sensory signals, such as proprioception, can be difficult. To begin to investigate these sensorimotor contributions, we recorded from flies whose legs had been removed in an attempt to impair naturalistic leg-based proprioceptive activity patterns such as those experienced during walking. Under these conditions, we observed large amplitude GLNO activity associated with movements of the abdomen, even in complete darkness (**Figure 3 S1**). On average, leftward abdomen bends were associated with activation of the left GLNO neurons and rightward abdomen bends with activation of the right GLNO neurons, consistent with the observation that abdomen bends and turns are correlated during tethered flight (Fenk et al., 2021). However, the correlation between abdomen bends and GLNO activity was quite variable, including periods where the average relationship described above reversed. In addition, most abdomen bends were not associated with appreciable GLNO activity. Together, these observations suggest a partial dissociation between GLNO activity and the fly’s proprioceptive state, raising the possibility of a contribution from an efference copy. For now, we refer to this signal as a “motor signal” to include potential contributions from motor efference and proprioception, a topic we return to in Discussion.

### GLNO neurons can encode rotational velocity using optic flow information

Every time an animal turns, the visual scene appears to sweep across the eye in the opposite direction, generating a reafferent optic flow signal that animals can use to assess whether their intended turn had the expected sensory consequences. In other situations, an externally applied force, such as a gust of wind, may cause an animal to rotate, and animals could use optic flow to detect such events and update their HD representation accordingly. With these considerations in mind, we next tested whether GLNO neurons are sensitive to optic flow, a possibility suggested by previous work on PEN neurons (Green et al., 2017) and other LNO neuron types (Currier et al., 2020; Lu et al., 2022; Lyu et al., 2022; Stone et al., 2017). We began by moving a 45-degree square wave grating in the leftward or rightward directions at 45 °/s for 5 seconds while recording from GLNO neurons in flies whose legs had been removed to reduce spontaneous motor signals. Consistent with the presence of an optic flow signal, the left GLNO neurons, which are active during left turns, are also driven by optic flow stimuli that would indicate to the fly that it was turning left. Similarly, the right GLNO neurons, which are active during right turns, are also driven by optic flow stimuli that would indicate to the fly it was turning right (**Figure 3G**). That is, the left and right GLNO neurons receive an optic flow signal that is tuned to the same direction as their motor input. GLNO neurons were largely insensitive to optic flow moving in their non-preferred direction (**Figure 3H**).

To further investigate the properties of the optic flow signal, we adopted an approach employed previously to parametrically characterize optic flow responses to a range of stimuli (Creamer et al., 2018). In particular, we presented flies with square wave gratings with different spatial wavelengths, from 15 degrees to 120 degrees, moving at a number of different temporal frequencies, from 1/16 Hz to 16 Hz (**Figure 3 S2**). When presented with visual motion in their preferred direction, GLNO neurons respond with a transient activation and variable habituation dynamics whose amplitudes depend on the stimulus’s temporal frequency and spatial wavelength (**Figure 3 S2 A**). Slow stimuli produced a small amplitude response that attenuated over the 5 second stimulus window. In contrast, stimuli moving at intermediate speeds (∼50 °/s) generated medium to large amplitude responses that attenuated very little. Stimuli moving at high speeds generated a large amplitude response that strongly attenuated over the course of 5 seconds. As a result, in the first ∼1.6 seconds after stimulus onset, GLNO neurons increase their activity in proportion to the stimulus’s rotational velocity, as expected for neurons encoding rotational velocity, but their activity peaks at velocities of ∼100 °/s and declines thereafter (**Figure 3 S2 C-E**). A similar trend holds for later time windows, but habituation lowers the peak velocity response to ∼50 °/s. We return to these results in Discussion when considering how flies may utilize optic flow to update their HD representation.

### Motor information overrides congruent optic flow signals

The results described above indicate that GLNO neurons can encode the fly’s rotational velocity using both motor and visual information. How do the GLNO neurons utilize these cues when both are present? To begin to address this question, we next recorded GLNO activity as flies with intact legs walked in closed-loop with a square wave grating, but while systematically varying the closed-loop gain from -2 to 2. With a closed-loop gain of 1, the visual scene moves in lockstep with the fly’s turns. With a closed-loop gain of 2, the visual scene moves twice as much as expected from the fly’s turns, but in the direction expected from the fly’s movement. And a closed-loop gain of - 1 causes the visual scene to move in the opposite direction than might be expected from the fly’s turn. Using this approach, we recorded GLNO activity with closed-loop gains from -2 to 2 in increments of 0.25.

During positive gain trials, a turn in one direction will generate congruent motor and optic flow signals that could together drive GLNO neurons in the same hemisphere (**Figure 4A**, top panel). We reasoned that if GLNO neurons combined motor and visual cues to estimate rotational velocity, then the GLNO tuning curves should systematically change with the closed-loop gain (**Figure 4B**, top panel). In contrast, if flies rely primarily on motor information at the expense of optic flow, then GLNO tuning curves should remain relatively invariant across all positive closed loop gains (**Figure 4B**, top panel). **Figure 4A** shows data from two trials: one with a closed loop gain of 0.25 and one with a closed loop gain of 2, plotted on the same axis for direct comparison. Even though the visual motion varied by a factor of 8 between these two trials, the difference in GLNO activity remained properly scaled to the fly’s turns, producing near-identical tuning curves (**Figure 4B**). Consistent with this, across all positive closed-loop gains, the slope of the GLNO tuning curve remained relatively invariant, with consistently high correlation coefficients (**Figure 4C**). These results suggest that GLNO neurons primarily rely on motor signals at the expense of optic flow cues even when they are congruent with the fly’s turns (**Figure 4C**, top schematic), effectively forgoing visuomotor integration in this context.

**Figure 4:**
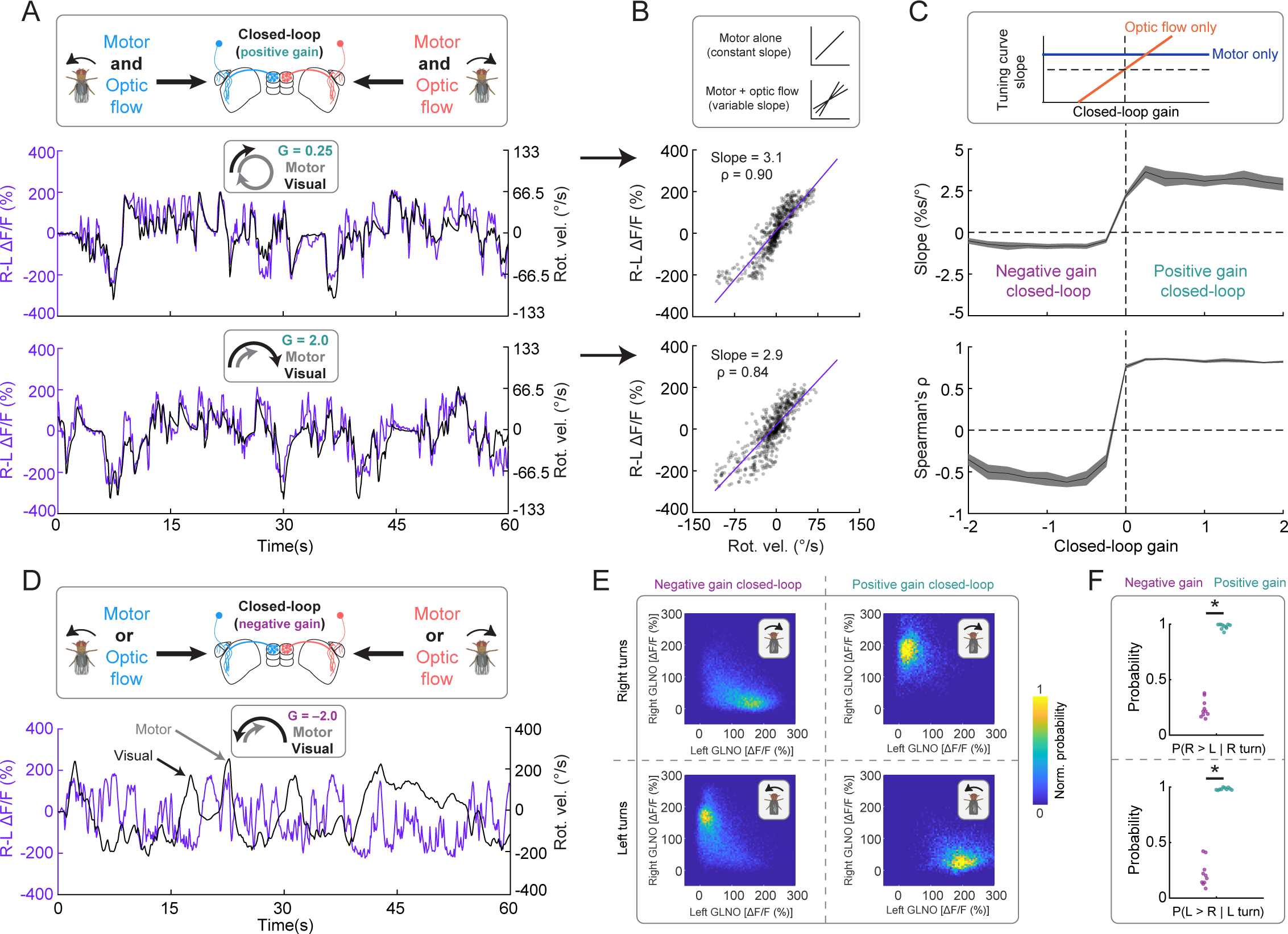
Sensorimotor competition between the left and right GLNO neurons. **A)** Top: illustration showing that during closed-loop experiments with different positive gains (from 0.25 to 2), a turn in one direction will generate a motor signal and a congruent optic flow signal whose magnitude scales with the closed-loop gain. Middle: example trial show- ing the difference in GLNO activity (right minus left, shown in purple) plotted alongside the fly’s rotational velocity (shown in black) when the closed-loop gain was set to 0.25, causing the visual scene to move 25% slower than the fly would expect based on the magnitude of its turn. Bottom: same as above, but when the closed-loop gain was set to 2, causing the scene to move twice as much as the fly would expect. Note that the two example trials are plotted on the same axis ranges for direct comparison. **B)** Top: schematic showing two competing models for how the GLNO tuning curves should change with the closed loop gain. If the GLNO neurons are only sensitive to the motor signal, the tuning curves should not change their shape when the closed-loop gain is altered (across positive values). Alternatively, if the GLNO neurons combine the motor and optic flow estimates, we would expect to see their tuning curves get steeper with increased closed-loop gains. Middle: scatter plot showing the GLNO tuning curve for the corresponding trial from (A). Bottom: same as above, but for the bottom trial from (A). Note that the tuning curves remain nearly identical even though the magnitude of the optic flow signal varies by a factor of 8 between the two trials. **C)** Fly-averaged (+/- s.e.m.) tuning curve slopes (top plot) and correlation coefficients (Spearman’s ρ) across all closed-loop gains, from -2 to 2 in steps of 0.25. The schematic at the top shows the predicted results from two competing models: one where the GLNO neurons only rely on motor information (blue curve) and one where they rely only on optic flow information (red curve). **D)** Top: illustration showing that during negative gain closed-loop experiments, a turn in one direction will generate a corresponding motor signal along with an optic flow signal indicating that fly turned in the opposite direction, setting up a sensorimotor conflict. Bottom: example trial showing the difference in GLNO activity plotted alongside the fly’s rotational velocity. The black arrow marks a turn where GLNO activi- ty followed the visual input. The gray arrow marks a turn where GLNO activity followed the motor signal. **E)** 2D histograms showing the normalized probability of the left (x-axes) and right (y-axes) GLNO activity across all negative gain (first column) and positive gain trials (second column) for both right turns (top panels) and left turns (bottom panels). Note localized regions of higher density in the state space. **F)** Top: beeswarm plot showing the probability that the right GLNO neurons were more active than the left GLNO neurons during a right turn across all flies (circles) for both negative gain (magenta) and positive gain (light blue) trials. Bottom: same as above but for left turns. GLNO neurons driven by the motor signal had a significantly higher probability of being active during positive gain trials relative to negative gain trials, as marked by the bars with asterisks, for both rightward (top panel) and leftward (bottom panel) turns (assessed with a Wilcoxon rank sum test with multiple comparison correction using the false discovery rate).

### Flip-flop dynamics may resolve visuomotor conflicts

We next asked whether the precedence of motor signals is maintained during negative gain trials, in which visual input is counter to that expected from the fly’s turns. While such contradictory signals are likely rare, they could occur in specific situations, such as when objects near the fly— other flies or blades of grass, for example—happen to move in the same direction as the fly is turning, generating incongruent optic flow. In situations like these, turns in one direction convey a motor signal to GLNO neurons in one hemisphere and an optic flow signal to the GLNO neurons in the contralateral hemisphere, producing a visuomotor conflict that requires resolution (**Figure 4D**, top schematic). Unlike positive gain trials, GLNO activity during negative gain trials is poorly correlated with the fly’s rotational velocity (**Figure 4C**). Instead, GLNO activity switches between following the motor input (gray arrow in **Figure 4D**) or the optic flow input (black arrow in **Figure 4D**). That is, the GLNO neurons appear to resolve the visuomotor conflict by relying on either the motor signal or the optic flow signal but not averaging or even integrating them. Consistent with this, when we looked at the probability of the left and right GLNO neurons occupying portions of their two-dimensional state space, we found that under negative gain conditions either the left or right GLNO neurons could be active during left or right turns, but the hemisphere receiving the optic flow signal was active more often (**Figure 4E-F**). Since the left and right GLNO neurons lack direct connections with each other, these results suggest that GLNO flip-flop dynamics are produced through reciprocal inhibition between their upstream inputs. Thus, lateral inhibition may function to resolve visuomotor conflicts by ensuring that GLNO neurons are driven by one cue at the expense of the other, though it remains unclear why a particular cue wins out at any given time. Together, these closed-loop experiments indicate that GLNO neurons rely on motor information in most circumstances, but can switch to relying on optic flow.

#### GLNO open-loop optic flow responses are consistent with lateral inhibition between hemispheres

How do GLNO neurons respond when the fly initiates a course-stabilizing turn to counter externally induced optic flow, as might be experienced on a windy day? To examine this question, we recorded GLNO activity while flies walked in open-loop with an optic flow stimulus previously found to generated large, sustained GLNO responses (a 45-degree square wave grating moving at 50 deg/s). Under these conditions, flies make frequent turns in the direction of visual motion in an attempt to correct for the perceived externally-induced rotation, a well-established behavior known as the optomotor response. During such turns, motor information will drive GLNO neurons in one hemisphere while visual information should drive GLNO neurons in the opposite hemisphere (**Figure 5A**, top panel). In contrast, during brief periods of immobility, only the optic flow cue will be present. Consistent with this, we found that the GLNO neurons driven by the optic flow stimulus were tonically activity during immobility (**Figure 5A**, blue circles). When flies initiated an optomotor response, two events occurred. First, the GLNO neurons driven by the motor signal turned on. Second—as might be expected from our earlier results (**Figure 4**)—the GLNO neurons that were receiving optic flow input turned off. This process could be reliably observed on a turn-by-turn basis (**Figure 5A**, red circles). In agreement with this, when we triggered GLNO activity on turns in the left or right direction, we found a large decrease in activity for GLNO neurons that were being driven by the optic flow input, results consistent across all 10 flies (**Figure 5B-C**). These results suggest that lateral inhibition prioritizes motor-signals generated during course-correcting turns over optic flow signals that would result from external perturbations (**Figure 5B**, top panels), and that optic flow signals may be primarily utilized when the motor signal is absent. Together, these results provide evidence for a circuit that facilitates visuomotor competition, where flies update their compass according to only one type of self-motion signal at a time (**Figure 6**).

**Figure 5:**
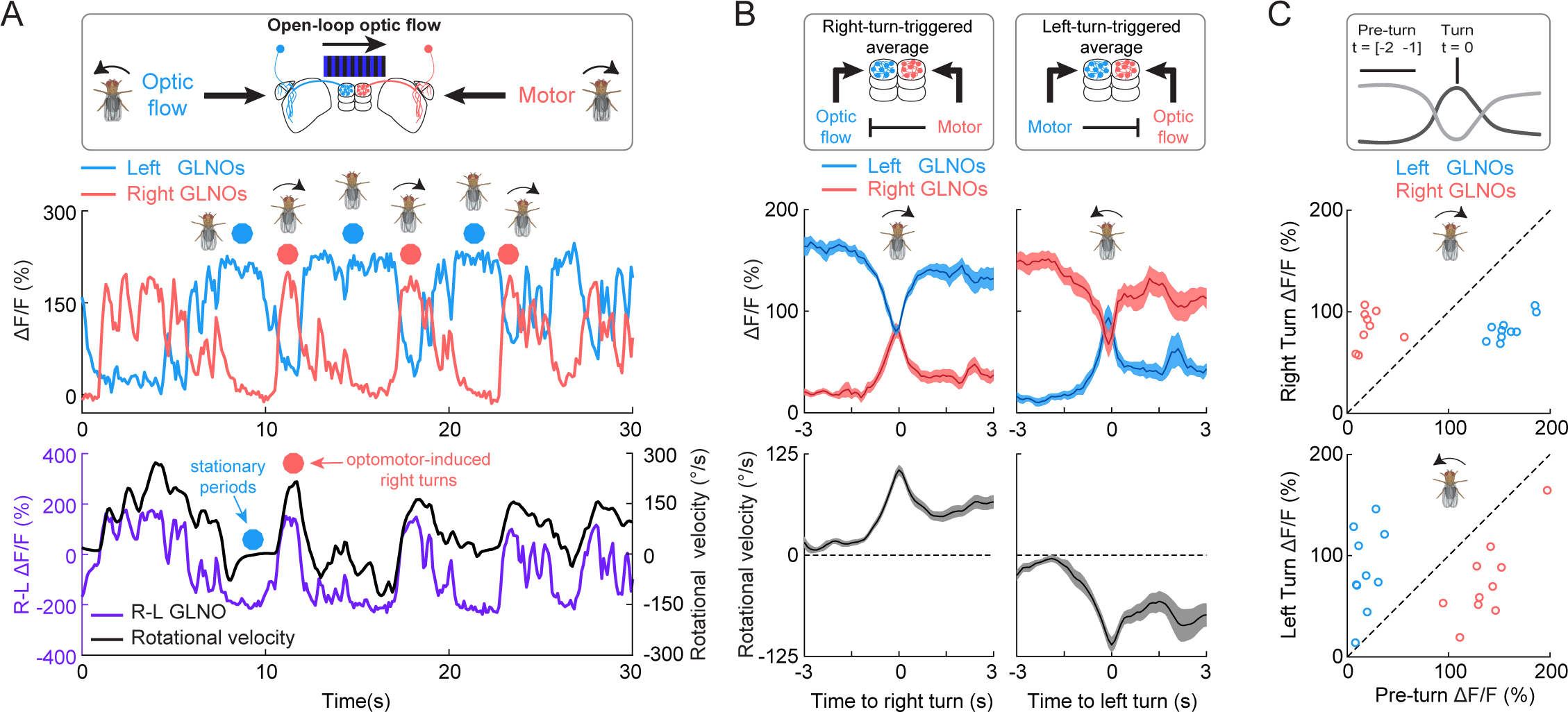
Reciprocal inhibition between left and right GLNO activity. **A)** Top: illustration showing that during open-loop experiments, optic flow will continually drive one GLNO population (left, in this case) while occasional optomotor responses will generate a motor signal in the other GLNO population (right, in this case). Middle: example trial showing calcium activity for the left (blue) and right (red) GLNO neurons during an open-loop optic flow experiment. Blue circles mark periods where the fly was largely stationary. Red circles mark three consecutive optomotor responses, when the fly makes a right turn. Notice that when the motor signal comes on (red trace), the GLNO neurons driven by optic flow (blue trace) get inhibited. Bottom: the difference in GLNO activity (right minus left, shown in purple) is plotted alongside the fly’s rotational velocity (shown in black). **B)** Turn-triggered average of GLNO activity for optomotor-induced turns in the rightward (left column) or leftward (right column) direc- tion. Top plots show the fly-averaged activity (+/- s.e.m.) in the left (blue) and right (red) GLNO neurons. Bottom plots show the fly averaged (+/- s.e.m.) rotational velocity. Turns were detected as peaks in the fly’s rotational velocity that exceeded +/- 40 °/s and that were preceded by a period (-3 to -1 s) of low rotational velocity (absolute average less than 40 °/s). Schematics at the top illustrate how these dynamics could be explained by an inhibitory connection between the upstream inputs to the left and right GLNO neurons. **C)** Scatter plots showing the per-fly GLNO activity before the turn (x-axis) compared to during the turn (y-axis), for turns in the rightward (top) or leftward (bottom) directions. All 10 flies showed a decrease in optic flow driven activity during both rightward (blue dots in top panel) and leftward (red dots in bottom panel) turns, leading to a significant difference in group medians as assessed by a Wilcoxon rank sum test with multiple comparison correction using the false discovery rate. Similarly, flies showed a significant increase in motor driven activity during both rightward (red dots in top panel) and leftward turns (blue dots in bottom panel).

**Figure 6:**
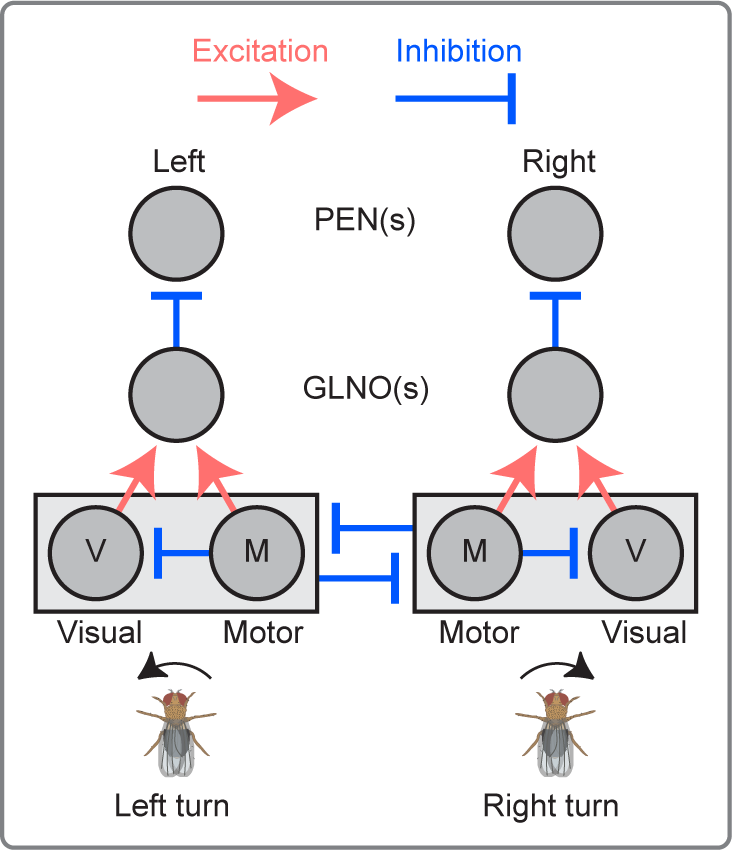
A conceptual circuit model that can account for GLNO sensorimotor dynamics. Schematic showing a functional architecture that can account for GLNO sensorimotor compu- tations, as described in the corresponding Discussion text. Connectomic tracing of GLNO inputs back to their sensory and motor sources proved difficult based on connectivity alone (Hulse et al., 2021), but the proposed functional architecture will help guide future physiological investigation. The left GLNO neurons, which are active during left turns, receive both visual (V) and motor (M) inputs that signal left turns, and vice versa for the right GLNO neurons. Within each hemisphere, the motor signal inhibits the optic flow signal, which can explain why the GLNO tuning curves change very little across all positive closed-loop gains (Figure 4). In addition, there is reciprocal inhibition between the left and right GLNO inputs, which could explain the GLNO flip-flop activity dynamics as well as the sensorimotor competition observed in our open loop (Figure 5) and negative closed-loop gain experiments (Figure 4). Note that other, similar architectures can also account for the above observations. For example, here we schematize the visual and motor inputs to GLNO neurons as being contained in a larger rectan- gle to highlight the possibility that the rotational velocity signal may come preformatted, though this need not be the case, an issue that requires future physiological investigation

## Discussion

Through a combination of whole-brain connectomics, causal circuit perturbations, and two-photon calcium imaging of neural activity during visuomotor behavior, our work identifies GLNO neurons as the main conduit of rotational velocity information to the fly HD network. While previous studies have shown that PEN neurons are conjunctively tuned to the fly’s HD and rotational velocity (Green et al., 2017; Turner-Evans et al., 2017), the source and sensorimotor nature of the rotational velocity signal was unknown. Our connectomic analysis reveals that GLNO neurons are the only PEN input that targets the left and right populations separately, a necessary requirement for producing EPG bump updates through the differential control of the left and right PEN bump amplitudes (**Figure 1B**). Consistent with this, reducing synaptic transmission from GLNO neurons causes the HD bump to uncouple from the fly’s actual HD (**Figure 2**). Two-photon calcium imaging of GLNO activity *in vivo* revealed that GLNO neurons can construct a rotational velocity signal using either motor or optic flow information (**Figure 3**). Under most circumstances, when visual and motor signals convey largely congruent velocity information, GLNO neurons rely on motor information at the expense of optic flow (positive gain experiments in **Figure 4**). However, when visual and motor signals convey opposing velocity estimates **(Figure 4C-F**), GLNO neurons toggle between following visual or motor information on a turn-by-turn basis, providing evidence for reciprocal inhibition between these upstream inputs to GLNO neurons. During open-loop experiments, optic flow signals were present during immobility, when the motor signal was absent, but were quickly attenuated when the fly initiated a turn (**Figure 5**), hinting at a situation-specific preference for particular cues.

### A simple conceptual model explains GLNO visuomotor processing

**Figure 6** shows a simple conceptual model that can account for our main experimental results. As described above (Figure 1B-C), GLNO neurons innervating the left LAL project to the left NO, where they inhibit the PEN neurons that innervate the right bridge; and GLNO neurons innervating the right LAL project to the right NO, where they inhibit the PEN neurons that innervate the left bridge (Franconville et al., 2018; Wolff et al., 2015). This allows the GLNO neurons to differentially control the left and right PEN bump amplitudes to drive compass updates (Green et al., 2017; Turner-Evans et al., 2017). Previous work identified this LAL-to-NO pathway, which is composed of distinct LNO neuron types, as a major source of the central complex’s self-motion information (Lu et al., 2022; Lyu et al., 2022; Stone et al., 2017), but tracing the LNO inputs back to their sensory and motor sources proved difficult (Hulse et al., 2021). Nevertheless, our physiological observations allow us to propose a functional architecture that can be used to guide future anatomical and physiological studies. As shown in Figure 6, GLNO neurons receive congruent motor and visual input conveying either leftward or rightward rotational velocity information. We schematize these visual and motor signals as contained within a larger rectangle to highlight the possibility that GLNO neurons may inherit their visuomotor estimate of rotational velocity from a single upstream neuron type since this signal may be relevant for other circuits and computations as well. One potential candidate is the PS196 neurons, which provide the largest input to GLNO neurons, though GLNO receive other LAL inputs as well, including from LAL139 and LAL184 (Hulse et al., 2021). To account for our observation that GLNO neurons rely on motor information at the expense of optic flow across all positive closed-loop gains (Figure 4), the motor signal likely inhibits congruent optic flow information on each side of the brain. Interestingly, previous studies have found evidence for motor signals inhibiting reafferent visual feedback in a widely studied class of visual motion-sensitive neurons known as lobula plate tangential cells (LTPCs; (Fenk et al., 2021; Fujiwara et al., 2022; Fujiwara et al., 2017; Kim et al., 2017a; Kim et al., 2015)). While the exact function of this inhibition in controlling the head and/or body direction during course stabilization is still being worked out, it raises the possibility that motor information may inhibit optic flow signals well upstream of the GLNO neurons. Lastly, to account for GLNO flip-flop dynamics (Figures 2-5) as well as visumotor competition during negative gain closed-loop experiments (Figure 4) and open-loop experiments (Figure 5), we suggest that GLNO inputs encoding leftward and rightward velocity reciprocally inhibit one another, a motif thought to be common in the LAL (Namiki and Kanzaki, 2016a). Together, the combination of motor signals that inhibit congruent optic flow and reciprocal inhibition between the left and right rotational velocity signals can account for our GLNO physiology data. Future work is required to map the functional architecture proposed here onto the neural circuits acting upstream of GLNO neurons (Erginkaya et al., 2023).

### A hierarchy of sensory and motor cues update the HD representation

Along with previous studies (Fisher et al., 2019; Green et al., 2017; Haberkern et al., 2022; Hulse et al., 2021; Kim et al., 2019; Okubo et al., 2020; Seelig and Jayaraman, 2015; Sun et al., 2017; Turner-Evans et al., 2017; Turner-Evans et al., 2020), our results indicate that flies update their HD representation using both sensory and motor cues. Like mammals (Savelli and Knierim, 2019; Taube, 2007; Taube et al., 1990a, b), flies update their compass using two processes that operate in parallel: integration of rotational velocity and drift correction, which tethers the HD bump to prominent directional cues like those from the sky (Haberkern et al., 2022; Hulse and Jayaraman, 2019). Directional sensory information is conveyed to the compass by distinct populations of ‘ring neurons’ that are tuned to a variety of cues, such as polarized light (Hardcastle et al., 2020), prominent visual landmarks (Seelig and Jayaraman, 2013), and the wind (Okubo et al., 2020). Through anti-Hebbian plasticity, ring neurons construct a map that tethers the HD bump’s position to the orientation of these directional sensory signals (Fisher et al., 2019; Fisher et al., 2022; Kim, 2021; Kim et al., 2019; Wilson, 2023). An important feature of this plasticity processes is that the ring neuron map should be strongest in the presence of highly reliable directional sensory information (Haberkern et al., 2022; Kim et al., 2019). Indeed, the EPG bump is reliably updated during closed-loop experience with a prominent visual landmark, including across a range of closed-loop gains (Seelig and Jayaraman, 2015). In a particularly striking example of visual input dictating the bump’s position, a previous study was able to form an inverted map using optogenetics, causing the HD bump to tether to a visual landmark that moved in the opposite direction than expected from the fly’s turns (Kim et al., 2019). These data suggest that a sufficiently strong directional map, when present, can largely determine the HD bump’s position, effectively overriding the contribution of angular path integration, at least on short time scales. Therefore, strong directional sensory information, when present, sits near the top of the sensorimotor hierarchy that updates the fly’s HD representation.

Particularly in the absence of a sufficiently strong directional map, flies must rely only on the integration of rotational velocity to update their compass. Our work indicates that GLNO neurons construct this velocity signal by relying on motor information at the expense of optic flow under most circumstances, including across all positive closed-loop gains (Figure 4). This suggests that flies may rely on optic flow information only in the worst-case scenario: when directional cues are absent and when motor efference signals indicate the fly is not turning in a particular direction, but the optic flow signal indicates otherwise. This may be particularly relevant during long-range flights (Leitch et al., 2021) that require flies to maintain a fixed heading for prolonged periods of time, an ability that is thought to rely on central complex circuitry (Giraldo et al., 2018; Green et al., 2019; Haberkern et al., 2022; Turner-Evans et al., 2020). During such relatively straight trajectories, the GLNO motor efference signal would be largely inactive. Yet, external perturbations, like a gust of wind, could transiently blow the fly off course. In worst case situations like these, the fly would have to rely on optic flow or proprioceptive feedback from the halters to update its compass and correct its course. If this is the case, it seems likely that such external perturbations would be relatively short in duration, suggesting that the GLNO neurons’ response to long-duration visual motion may be less relevant than their tuning during the first ∼1 second of visual motion, when they function as relatively accurate visual motion estimators up to ∼100 °/s (Figure 3 **S2**). Indeed, the fact that GLNO neuron show strong habituation to fast, prolonged visual motion could reflect a hard-coded structural prior biasing the system against such cues. Overall, these considerations indicate that flies update their compass using a hierarchy of cues, with strong directional sensory information at the top, motor efference signals in the middle, and optic flow and/or proprioception at the bottom.

Why might flies exclusively rely on motor efference at the expense of congruent optic flow that might, under ideal conditions, improve their estimate of rotational velocity? One possibility is that motor efference copies are more reliable in most ethological circumstances. Flies may have evolved powerful mechanisms to calibrate their efference copies, whereas constructing an absolute estimate of rotational velocity from optic flow alone is algorithmically challenging, requiring machinery to normalize for moment-to-moment changes in the statistics of the visual scene, such as contrast and spatial wavelength. Even so, certain motor contexts can give rise to rotational optic flow even if the fly isn’t turning, such as when another object passes by one side of the fly. Similarly, flies walking through 3D scenes may encounter situations that generate rotational optic flow, such as walking past a patch of grass, even if the fly isn’t turning (Haberkern et al., 2022). Another possibility is that relying on a single cue bypasses the need for complex visuomotor integration computations. For example, if the fly were to combine visual and motor estimates, then it would need a mechanism to scale the motor signal when the visual signal is absent, such as when walking in darkness.

### Velocity estimation in mammals and invertebrates

How might these invertebrate results relate to the extensive work on self-motion perception in mammals (reviewed in (Cullen, 2019; Cullen and Taube, 2017; Laurens and Angelaki, 2018; Noel and Angelaki, 2022))? Self-motion signals can be found throughout the mammalian brain, including cortical areas composing the dorsal visual stream (Britten, 2008) and subcortical structures like the vestibular nuclei (Cullen, 2019). How these various self-motion pathways update internal representations supporting navigation behaviors remains unknown, but vestibular pathways are thought to play a prominent role in updating the mammalian HD system (Clark and Taube, 2012; Cullen and Taube, 2017; Hulse and Jayaraman, 2019; Yoder and Taube, 2014). In particular, several studies have found that impairing vestibular function abolishes accurate angular path integration by HD cells (Butler et al., 2017; Muir et al., 2009; Stackman et al., 2002; Stackman and Taube, 1997; Valerio and Taube, 2016; Yoder and Taube, 2009). Interestingly, these manipulations often cause HD cells to untether from sensorimotor input and fire bursts of activity, a pattern consistent with the HD bump drifting along its ring manifold (Cullen and Taube, 2017). These observations resemble the EPG bump drift we observed after reducing synaptic transmission from the GLNO neurons (Figure 2), though in both cases it remains unclear which forces are causing the drift. While mammals require an intact vestibular system to update their HD representation, the upstream vestibular inputs, originating in the vestibular nuclei, are known to be suppressed during active movements (Carriot et al., 2013; Carriot et al., 2015; Medrea and Cullen, 2013; Roy and Cullen, 2001, 2004), suggesting that vestibular pathways preferentially transmit rotational information generated by passive or unexpected rotations. Similarly, LPTC neurons in *Drosophila*, a core component of early motion vision pathways, are known to receive motor signals that can cancel reafferent optic flow signals, allowing visually-driven self-motion activity only during externally-induced rotations that may require gaze and/or course stabilization (Chiappe, 2023; Fenk et al., 2021; Fujiwara et al., 2022; Fujiwara et al., 2017; Kim et al., 2017a; Kim et al., 2015). Similarly, our results suggest that during active movements, flies rely on motor information rather than optic flow to update their HD representation (Figure 4; positive gain results). Thus, in both mammals and flies, it seems likely that the HD representation is updated by distinct sensorimotor signals depending on context.

A recent proposal suggests that the final rotational velocity signal updating the mammalian HD representation is the output of a Kalman filter (Laurens and Angelaki, 2017, 2018). According to this view, the system uses a forward model (Cullen and Taube, 2017; Webb, 2004) to cancel expected sensory reafference and integrates any resulting sensory prediction error with motor efference signals to form a multimodal estimate of rotational velocity (Brooks and Cullen, 2014, 2019; Cullen and Brooks, 2015; Laurens and Angelaki, 2018). In this framework, the inhibition of early vestibular or motion vision pathways is the result of sensory prediction signals, and any remaining activity in these pathways is interpreted as a sensory prediction error. While visual and motor signals are known to be important for updating the mammalian HD representation (Arleo et al., 2013; Blair and Sharp, 1996; Stackman et al., 2003; Yoder et al., 2011), it remains unclear where these signals originate or whether their usage is consistent with that of a Kalman filter. In contrast, our results suggest that under most circumstances, *Drosophila* rely on motor efference signals to construct a rotational velocity estimate, effectively bypassing the need for complex visuomotor integration machinery. Similarly, flies may only rely of optic flow information during unexpected rotations, when the motor signal would be absent. These results suggest that the fly’s compact neural circuits may rely on heuristics that were learned over evolutionary timescales and that may require less neural machinery to implement than optimal visuomotor integration.

## Future work

Our work also suggests several lines of future inquiry. First, the two PEN subtypes, PEN_a and PEN_b, have distinct bump dynamics (Green et al., 2017) and network connectivity patterns (Hulse et al., 2021; Turner-Evans et al., 2020), but how these two populations function together to update the compass remains poorly understood. GLNO neurons synapses onto both PEN subtypes, but distinct postsynaptic receptor composition could account for their characteristic turning responses (Green et al., 2017) and functional connectivity (Franconville et al., 2018). Second, while our works suggests that GLNO neurons rely on motor information at the expense of optic flow in most circumstances, it remains possible that this balance could be different in flying flies, when motor efference copies may be less reliable. It is also possible that sufficiently bright optic flow stimuli, which are difficult to use in two-photon calcium imaging experiments, could favor the use of optic flow in some circumstances. In addition, the contribution of proprioceptive feedback from the halteres remains unexplored (but see (Kathman and Fox, 2019)), largely due to the requirement to record from tethered flies using existing technology. Similarly, how GLNO neurons function in complex 3D scenes is unknown (Haberkern et al., 2022). And finally, interpretating the potential function of the GLNO flip-flop dynamics could be aided by membrane potential recordings, which are challenging due to the location of the GLNO soma. If GLNO neurons have a resting firing rate, reciprocal inhibition could serve a ‘seesaw’ function by reducing the activity of left GLNO neurons as right GLNO neurons increase their activity (or vice versa), perhaps to ensure the difference in the left and right PEN bumps remains symmetric about their mean. On the other hand, if GLNO neurons are silent at rest, reciprocal inhibition could primarily function to inhibit incongruent velocity signals, ensuring that whichever cue is strongest wins out. Next-generation voltage indicators may help address this issue and allow for the study of sensorimotor competition at faster timescales.

## Outlook

Flexible navigation requires animals to construct robust self-motion estimates that can accurately update internal representations across different sensorimotor contexts. Our work identifies a self-motion pathway used to update the fly’s HD network, and physiological results indicate that flies primary rely on motor information at the expense of optic flow. More broadly, given that HD networks have been described in a variety of phylogenetically distant animals, it may now be possible to dissect how distinct species construct velocity estimates that are tailored to their unique sensorimotor experiences and navigational needs. While all animals must rely on sensory and motor information to construct these velocity estimates, working out the implementation level details has proven challenging, especially in vertebrates. Complementing detailed circuit studies in flies, future connectomic and physiology studies in small vertebrate species, such as larval zebrafish (Petrucco et al., 2022; Yang et al., 2022), may provide a path towards a deeper understanding of self-motion computations across phylogeny.

## Acknowledgements

We thank Vasily Goncharov, Chris McRaven, and Tim Hanson for microscopy support; Steve Sawtelle for treadmill assistance; Michael Reiser, Matthew Isaacson, and Jinyang Liu for sharing their G4 panel display system; Bill Biddle, Jon Arnold, and Tobias Goulet for fly pyramid fabrication; Will Dickson for BIAS support; and Ann Hermundstad, Sandro Romani, and members of the Jayaraman lab for many helpful discussions and feedback on the manuscript.

## Materials and methods

### Fly stocks

All fly stocks were maintained on cornmeal-based fly food at 23°C and 60% relative humidity with a 16 hour light period (6am to 10pm) and an 8 hour dark period (10pm to 6am), with all experiments occurring during the light period. **Table 1** contains a complete list of genotypes used for all experiments. Gen1 GAL4 and LexA drivers targeting the GLNO neurons (GMR76e11) and EPG neurons (VT025957; GMR27f02) were generated by (Jenett et al., 2012) and (Tirian and Dickson, 2017). Split-GAL4 drivers (Pfeiffer et al., 2010) targeting the GLNO neurons (SS46517 and SS46524) were generated by (Wolff and Rubin, 2018). GCaMP6 flies were generated by (Chen et al., 2013); GCaMP7 flies were generated by (Dana et al., 2019); and flies used in Shibire^ts^ experiments were obtained from (Turner-Evans et al., 2020). **Table 2** contains a list of all constructs present in each fly genotype, as well as their source.

**Table 1:**
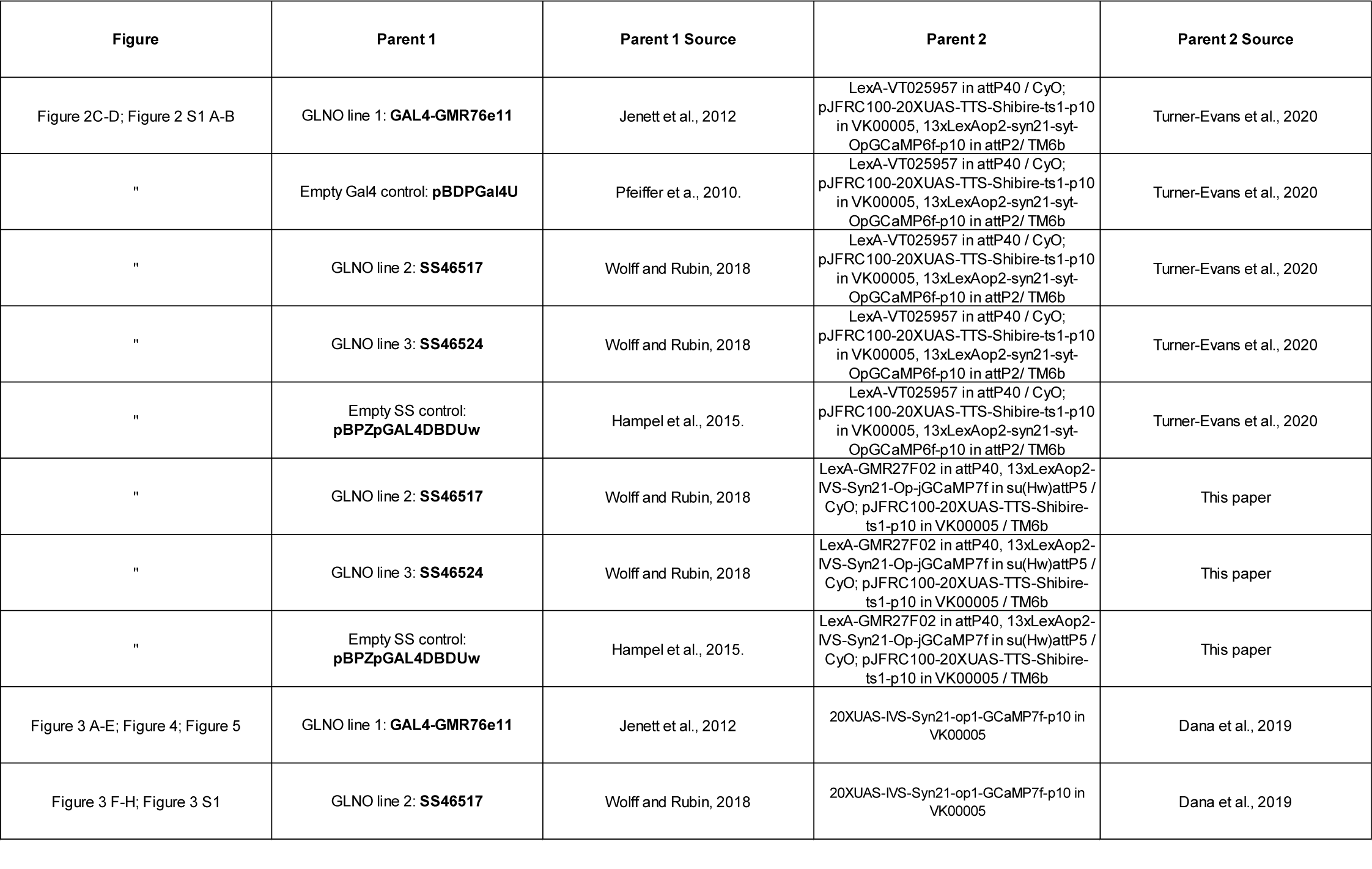
Genotypes for associated figures.

| Figure | Parent 1 | Parent 1 Source | Parent 2 | Parent 2 Source |
| --- | --- | --- | --- | --- |
| Figure 2C-D; Figure 2 S1 A-B | GLNO line 1: **GAL4-GMR76e11** | Jenett et al., 2012 | LexA-VT025957 in attP40 / CyO; pJFRC100-20XUAS-TTS-Shibire-ts1-p10 in VK00005, 13xLexAop2-syn21-syt-OpGCaMP6f-p10 in attP2/ TM6b | Turner-Evans et al., 2020 |
| " | Empty Gal4 control: **pBDPGal4U** | Pfeiffer et a., 2010. | LexA-VT025957 in attP40 / CyO; pJFRC100-20XUAS-TTS-Shibire-ts1-p10 in VK00005, 13xLexAop2-syn21-syt-OpGCaMP6f-p10 in attP2/ TM6b | Turner-Evans et al., 2020 |
| " | GLNO line 2: **SS46517** | Wolff and Rubin, 2018 | LexA-VT025957 in attP40 / CyO; pJFRC100-20XUAS-TTS-Shibire-ts1-p10 in VK00005, 13xLexAop2-syn21-syt-OpGCaMP6f-p10 in attP2/ TM6b | Turner-Evans et al., 2020 |
| " | GLNO line 3: **SS46524** | Wolff and Rubin, 2018 | LexA-VT025957 in attP40 / CyO; pJFRC100-20XUAS-TTS-Shibire-ts1-p10 in VK00005, 13xLexAop2-syn21-syt-OpGCaMP6f-p10 in attP2/ TM6b | Turner-Evans et al., 2020 |
| " | Empty SS control: **pBPZpGAL4DBDUw** | Hampel et al., 2015. | LexA-VT025957 in attP40 / CyO; pJFRC100-20XUAS-TTS-Shibire-ts1-p10 in VK00005, 13xLexAop2-syn21-syt-OpGCaMP6f-p10 in attP2/ TM6b | Turner-Evans et al., 2020 |
| " | GLNO line 2: **SS46517** | Wolff and Rubin, 2018 | LexA-GMR27F02 in attP40, 13xLexAop2-IVS-Syn21-Op-jGCaMP7f in su(Hw)attP5 / CyO; pJFRC100-20XUAS-TTS-Shibire-ts1-p10 in VK00005 / TM6b | This paper |
| " | GLNO line 3: **SS46524** | Wolff and Rubin, 2018 | LexA-GMR27F02 in attP40, 13xLexAop2-IVS-Syn21-Op-jGCaMP7f in su(Hw)attP5 / CyO; pJFRC100-20XUAS-TTS-Shibire-ts1-p10 in VK00005 / TM6b | This paper |
| " | Empty SS control: **pBPZpGAL4DBDUw** | Hampel et al., 2015. | LexA-GMR27F02 in attP40, 13xLexAop2-IVS-Syn21-Op-jGCaMP7f in su(Hw)attP5 / CyO; pJFRC100-20XUAS-TTS-Shibire-ts1-p10 in VK00005 / TM6b | This paper |
| Figure 3 A-E; Figure 4; Figure 5 | GLNO line 1: **GAL4-GMR76e11** | Jenett et al., 2012 | 20XUAS-IVS-Syn21-op1-GCaMP7f-p10 in VK00005 | Dana et al., 2019 |
| Figure 3 F-H; Figure 3 S1 | GLNO line 2: **SS46517** | Wolff and Rubin, 2018 | 20XUAS-IVS-Syn21-op1-GCaMP7f-p10 in VK00005 | Dana et al., 2019 |

**Table 2:** Fly stocks and their source.

| Construct | Source |
| --- | --- |
| GAL4-GMR76e11 | Bloomington Stock Center (RRID:BDSC_39933) |
| pBDPGal4U | Bloomington Stock Center (RRID:BDSC_68384) |
| SS46517 | Bloomington Stock Center (RRID:BDSC_76018) |
| SS46524 | Bloomington Stock Center (RRID:BDSC_76018) |
| pBPZpGAL4DBDUw | Bloomington Stock Center (RRID:BDSC_79603) |
| LexA-GMR27F02 in attP40 | Bloomington Stock Center (RRID:BDSC_52748) |
| LexA-VT025957 in attP40 | Janelia (robot ID: 3010906) |
| 13xLexAop2-syn21-syt-OpGCaMP6f-p10 in attP2 | Bloomington Stock Center (RRID:BDSC_44277) |
| 13xLexAop2-IVS-Syn21-Op-jGCaMP7f in su(Hw)attP5 | Janelia (robot ID: 3041065) |
| 20XUAS-IVS-Syn21-op1-GCaMP7f-p10 in VK00005 | Bloomington Stock Center (RRID:BDSC_79031) |
| pJFRC100-20XUAS-TTS-Shibire-ts1-p10 in VK00005 | Janelia (robot ID: 1117405) |

### Fly preparation for imaging

Flies (females, age 5–9 d, *n* = 10 flies per group) were prepared for imaging as previously described (Seelig et al., 2010; Seelig and Jayaraman, 2015). Briefly, flies were anesthetized at 4 °C and their proboscis was immobilized with wax to reduce brain movements. Next, flies were tethered to a tungsten pin glued to their thorax (Loctite AA 3972), and their head/thorax was positioned and fixed to a recording chamber with light-cured adhesive (Fotoplast gel, Dreve). To gain optical access to the brain, a section of cuticle between the ocelli and antennae was removed, along with the underlying fat and air sacs. Throughout the experiment, the head was submerged in saline containing (in mM): NaCl (103), KCl (3), TES (5), trehalose (8), glucose (10), NaHCO3 (26), NaH2PO4 (1), CaCl2 (2.5) and MgCl2 (4), with a pH of 7.3 and an osmolarity of 280 mOsm.

### Spherical treadmill system and data acquisition

Following dissection, flies were positioned on an air-supported polyurethane foam ball (8 mm diameter, 47 mg) under the microscope and allowed to walk. Rotations of the ball were tracked at 500 Hz, as described previously (Seelig et al., 2010). Behavioral data and imaging timestamps were recorded using WaveSurfer (http://wavesurfer.janelia.org/). Side and rear views of the fly were recorded from two complementary metal oxide semiconductor cameras (Grasshopper3, FLIR) using BIAS (IO Rodeo). Flies were given 5-10 minutes to adjust to the ball before imaging began.

### Two-photon calcium imaging

Calcium imaging was performed with a custom-built two-photon microscope (MIMMS 2.0, https://www.janelia.org/open-science/mimms-21-2020) controlled with ScanImage 2020 (Vidrio Technologies; (Pologruto et al., 2003)). Excitation of genetically-encoded calcium indictors (GCaMP6f or GCaMP7f; (Chen et al., 2013; Dana et al., 2019)) was generated with an infrared (930 nm), femtosecond-pulsed (pulse width ∼ 110 fs) laser (Chameleon Discovery, Coherent) with 15-20 mW of power, as measured after the objective (60x Olympus LUMPlanFL/IR with NA=0.9 or 40x Olympus LUMPlanFLN with NA=0.8 or 20x Olympus XLUMPLFLN with NA =1.0). Fast *Z* stacks (8 planes with 3 to 6 µm spacing and three fly-back frames) were collected at 10 Hz by raster scanning (128 × 128 pixels; ∼25 × 25 µm^2^ or ∼50 x 50 µm^2^ for NO imaging and ∼100 × 100 µm^2^ for EB imaging) using an 8 kHz resonant-galvo system and piezo-controlled *Z* positioning. Focal planes were selected to cover the full extent of EPG processes in the EB or GLNO processes in the NO. Emitted light was directed (primary dichroic: 735; secondary dichroic: 594), filtered (filter A: 680 SP; filter B: 525/50), and detected with a GaAsP photo-multiplier tube (H10770PB-40, Hamamatsu).

### Visual display

The visual display system was constructed from an updated version (referred to as G4.0) of the light-emitting diode panels described in (Reiser and Dickinson, 2008). The visual arena was 22.5 cm in diameter and covered 240° in azimuth and ∼55° in elevation using a grid of 192 × 64 pixels (UV light-emitting diodes; emission peak: 375 nm; 2.5 mm pixel spacing). The display was refreshed synchronously at 500 Hz. The diameter of each pixel’s subtended area is at most 1.25° on the fly eye. To filter out green light emitted from the UV LEDs, we constructed a glass screen (Hoya B370) which strongly attenuated wavelengths from 500-700 nm. Further filtering was achieved through one Roscolux R59 gel filter and three Roscolux R39 and gel filters, along with a scattering material that reduced reflections (BrightWhite, BrightView Technologies). As in previous versions (Reiser and Dickinson, 2008), the angular position of a visual patterns (described below) was updated by feeding the display system a 0-10 V control signal. To update the visual pattern in a closed-loop fashion, we wrote a custom LabVIEW VI that converted the raw treadmill outputs, which encode the fly’s yaw velocity, into a 0-10 V signal encoding the fly’s virtual head direction, with a user-defined scaler that could be used to change the closed-loop gain (Figure 4). For open-loop control, the LabVIEW VI swept that 0-10 V range with a user-defined angular velocity (Figure 3 G-H; Figure 5).

### Recording EPG bump dynamics while reducing GLNO vesicle release

Experiments involving shibire^ts^ (Figure 2 and Figure 2 S1) were performed as described in (Turner-Evans et al., 2017; Turner-Evans et al., 2020). Briefly, shibire^ts^ is a temperature-sensitive mutant of the gene encoding *Drosophila*’s dynamin ortholog, a protein involved in vesicle endocytosis (Kitamoto, 2001). Shibire^ts^ functions normally at room temperature (∼22° C) but decreases vesicle endocytosis and therefore reduces synaptic release at elevated temperatures (31- 32° C). To test whether GLNO neurons are required for generating accurate EPG bump updates (Figure 2), we expressed shibire^ts^ in GLNO neurons using one of three drivers (76e11-Gal4, SS46517, or SS46524) with either GCaMP6f or GCaMP7f in EPG neurons using VT25957-LexA or 27F02-LexA. Control flies also expressed GCaMP in EPG neurons but the GLNO driver was swapped for an “empty” GAL4 (or split-GAL4) which lacks the enhancer element needed to drive expression (see **Table 1** for lines). To start, EPG dynamics were recorded at room temperature (∼22° C) during 6 minutes of darkness followed by 4 minutes where flies were given closed-loop control of a 20° bar. To deplete GLNO neurons of vesicles, the saline bath was then raised to 31- 32° C for 15 minutes while flies walked in closed loop with the 20° bar. Finally, EPG dynamics were assessed at the elevated temperature, during 4 minutes with the bar following by 6 minutes in darkness.

### Recording GLNO activity across different visuomotor contexts

Flies with GCaMP7f expressed in GLNO neurons (**Table 1**) were placed on the spherical treadmill and allowed to walk, first in darkness (Figure 3), then in closed-loop (Figure 4) and open-loop (Figure 5) with a square-wave grating shown to produced sustained optic flow responses in GLNO neurons (45° spatial wavelength; Figure 3). During open-loop trials, the square-wave grating moved at 50°/s in either the clockwise or counterclockwise direction. Across closed-loop trials, the gain was adjusted from -2 to 2 in increments of 0.25, presented in a pseudorandom order. Experiments involved 22 consecutive one-minute trials, starting with one trial of darkness (Figure 3), followed by 17 closed-loop trials, and 4 open loop trials (2 clockwise; 2 counterclockwise).

### Recording GLNO responses to optic flow in immobilized flies

To parametrically characterize GLNO optic flow tuning (Figure 3 G-H, Figure 3 **S2**), we presented immobilized flies with square-wave gratings with 6 different spatial wavelengths (in degrees: 22.5, 30, 45, 60, 90, 120) and 17 temporal frequencies (in Hz: 0.0625, 0.0938, 0.125, 0.1875, 0.25, 0.375, 0.5, 0.75, 1, 1.5, 2, 3, 4, 6, 8, 12, 16). Each square-wave grating of a particular spatial wavelength and temporal frequency was presented four times, twice in the clockwise direction and twice in the counterclockwise direction, with counterbalancing across trial blocks. Each grating was presented for 5 seconds with a 5 second inter-trial interval. Prior to imaging GLNO activity (**Table 1**), flies had their legs removed and stumps glued to reduce spontaneous motor activity.

### EPG calcium imaging analysis

Analysis of behavioral and imaging datasets were carried out using MATLAB (MathWorks). Circular data were analyzed using the CircStat toolbox (Berens, 2009). For EPG imaging experiments (Figure 2), each *Z*-stack was reduced to a single frame using a maximum intensity projection. An ellipse was manually drawn around the perimeter of the EB and automatically segmented into 32 equal-area, wedge-shaped regions of interest (ROIs). The number of ROIs was chosen to match the number of anatomically defined EB demi-wedges (Wolff et al., 2015). Activity within each ROI was averaged for each frame, producing 32 ROI time series. For each ROI time series, baseline fluorescence (*F*0) was defined as the average of the lowest 10% of samples. Δ*F*/*F* was computed as (*F* − *F*0)/*F*0 × 100, where *F* is the instantaneous fluorescence from the raw ROI time series. Each ROI’s Δ*F*/*F* time series was filtered with a third order Savitzky– Golay filter over seven frames (Seelig and Jayaraman, 2015). To compute the bump’s instantaneous position and strength, we used the population vector average (PVA; Figure 2B). For each frame, the PVA was computed by taking the circular mean of vectors whose angles were the ROI’s wedge positions and whose length was equal to the ROI’s Δ*F*/*F*. The magnitude of this mean resultant vector length was normalized to have a maximum possible length of one. As described previously (Seelig and Jayaraman, 2015), there is a fly-specific angular offset between the position of the bump in the EB and the position of the bar stimulus. This difference was subtracted for display purposes in Figure 2.

### GLNO calcium imaging analysis

Preprocessing of GLNO imaging data was performed similar to the EPG preprocessing described above: Z-stacks were reduced to a single frame using a maximum intensity projection, the left and right GLNO processes (in the NO) were used as regions of interest, activity within each ROI was averaged for each frame, and Δ*F*/*F* was computed. GLNO ROIs were automatically drawn using a multistep process: the collection of frames (128 x 128 x samples) was filtered with a 3D gaussian filter with a standard deviation of 2 pixels to smooth single-pixel noise; this collection of smoothed frames was reshaped into a matrix whose columns contained each pixel’s timeseries (samples x 128^2^); nonnegative matrix factorization was used to decompose the pixel timeseries matrix into two components whose spatial profiles mapped onto the left and right GLNO processes; finally, each component was scaled (dividing by the 98^th^ percentile across pixels) and a binary mask was created by thresholding above 0.8.

### Connectomic analysis of lateralized PEN inputs

In Figure 1D we analyze the hemibrain connectome (Hulse et al., 2021; Scheffer et al., 2020) to determine which neurons provide lateralized input to the PEN neurons, a necessary requirement for PEN-driven bump updates. For every neuron that provides input to a PEN_a neuron or a PEN_b neuron, we plotted the absolute difference in its total input to the right and left PEN populations. If a neuron provides equal input to the left and right PEN populations, this difference will be close to 0, while a highly lateralized input will yield a positive value that is far from 0. We restricted our analysis to neurons with at least 20 synapses, summed across all postsynaptic PEN neurons, and we excluded PEN_a, PEN_b, and EPG neurons as candidate presynaptic types. We accessed version 1.2.1 of the neuPrint database through its R API, neuprintr (http://natverse.org/neuprintr/; (Bates et al., 2019)), with additional postprocessing using the neuprintrExtra package (https://github.com/jayaraman-lab/neuprintrExtra; (Franconville, 2022)).

### Quantifying the correlation between EPG bump position and the fly’s virtual head direction

In Figure 2 and Figure 2 **S1** we analyze how the correlation between the fly’s virtual head direction and the angular position of the EPG bump in the EB is affected by reducing neurotransmitter release from the GLNO neurons. Each trial consisted of 6 minutes in darkness or 4 minutes in closed-loop with a 20° bar, both at room temperature and at the restrictive temperature (∼31° C). For each trial we broke the EPG imaging data into chunks where the PVA length was at least 0.075, which allowed for unambiguous tracking of the bump’s position. During periods of immobility, the EPG bump amplitude is reduced in amplitude (Seelig and Jayaraman, 2015), and our threshold on the PVA functioned to remove periods like these, where the bump could not be reliably tracked. We further restricted our analysis to chunks exceeding 7 seconds in length. For each of these chunks, we performed a robust linear regression between the fly’s unwrapped head direction and the unwrapped EPG bump position, yielding a correlation coefficient that captures the extent to which bump movements are driven by the fly’s turns. For trials that were broken into more than one chunk, the correlation coefficient was averaged across chunks. Similarly, for each trial we also computed the average bump amplitude and fly’s forward and rotational velocity (Figure 2 **S1 B-D**). To statistically assess whether reducing GLNO synaptic transmission affects the magnitude of these parameters, we compared group medians by performed a two-sided Wilcoxon rank sum tests between groups of flies expressing Shibire^ts^ in GLNO neurons and genetic controls, both at room temperature and at the restrictive temperature. We corrected for multiple comparisons by adjusting the significance level using the false discover rate (Benjamini and Hochberg, 1995), with an initial significance level of p=0.05. This statistical approach was used to compare group medians in Figure 4F and Figure 5C as well.

### Fitting sigmoidal and linear models to GLNO tuning curves

In Figure 3 A-D, we quantified how GLNO activity changes as a function of the fly’s rotational velocity as it walks in darkness for 60 seconds. As described in previous work (Turner-Evans et al., 2017), we convolved the fly’s raw rotational velocity signal with a kernel that mimics the dynamics of GCaMP6f, given by the following equation:

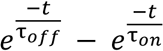

We used a kernel length of 5 seconds (0 ≤ t ≤ 5) with τ_off_ = 0.63 and τ_on_ = 0.13, normalized so the kernel summed to 0. Lowpass filtering the raw rotational velocity signal allowed us to compare it with GLNO calcium activity on a shared timescale. Constructing this kernel requires an estimate of the calcium sensor’s on and off time constants, which show considerable variability according to the number of action potentials and the neuron types they were measured in (Chen et al., 2013). Given these complexities, for consistency we use the same time constants as in (Turner-Evans et al., 2017), for both GCaMP6f and GCaMP7f recordings.

Following convolution, we plotted the difference in GLNO activity (right-left) as the function of the fly’s rotational velocity, as shown in Figure 3 **D**. As in previous work (Turner-Evans et al., 2017), we fit these tuning curves with a sigmoidal function given by:

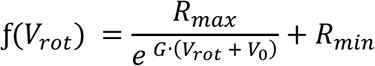

From these fits, we could compute the maximum tuning curve slope and the proportion of the variance explained by the model (Figure 3 **E**).

In Figure 4, we analyze how GLNO tuning curves change as a function the virtual reality system’s closed-loop gain. Under negative gain conditions, GLNO activity is no longer a sigmoidal function of the fly’s rotational velocity (Figure 4D). Instead, negative gain tuning curves were better fit by linear functions. Since linear regression also provides quality fits to positive gain trials (Figure 4B), we used this approach to estimate how the tuning curve slope and correlation coefficient varies with the closed-loop gain (Figure 4C).

### Correlation between GLNO activity in darkness and abdomen bends in legless flies

For the GLNO imaging experiments during spontaneous abdomen bends in darkness (Figure 3 **S1),** the flies were recorded synchronously from both side and rear views with two monochromatic cameras (Grasshopper3, FLIR, GS3-U3-23S6M-C) and the BIAS software (IO Rodeo). Frame acquisitions were externally triggered from a customized LabVIEW application (National Instruments) at 40Hz. Changes in abdomen posture were estimated from the rear-view frames using the Animal Part Tracker ((Kabra et al., 2022), https://github.com/kristinbranson/APT) by tracking the position of the tip of the abdomen relative to a static position on the dorsal thorax. From the raw abdomen tip positions, a moving median was calculated over a sliding window of 5000 samples along the vertical axis to estimate the non-biased posture at each abdomen bending angle. Horizontal deviations from this profile were then normalized to the overall standard deviation. Vertical tip positions were trend-corrected in time using a moving median with a 12000 sample sliding window and normalized to the moving 0.05-quantile. To calculate the correlations with the GLNO neuronal activity, which were recorded at approximately 10Hz, the abdomen tip positions were linearly interpolated at the GLNO neurons recorded time points. Discrete bouts of GLNO activity were generated by splitting the time series at segments where both left and right GLNO activities were low (<75%) for at least 10 frames.

---

**Figure 2 S1:**
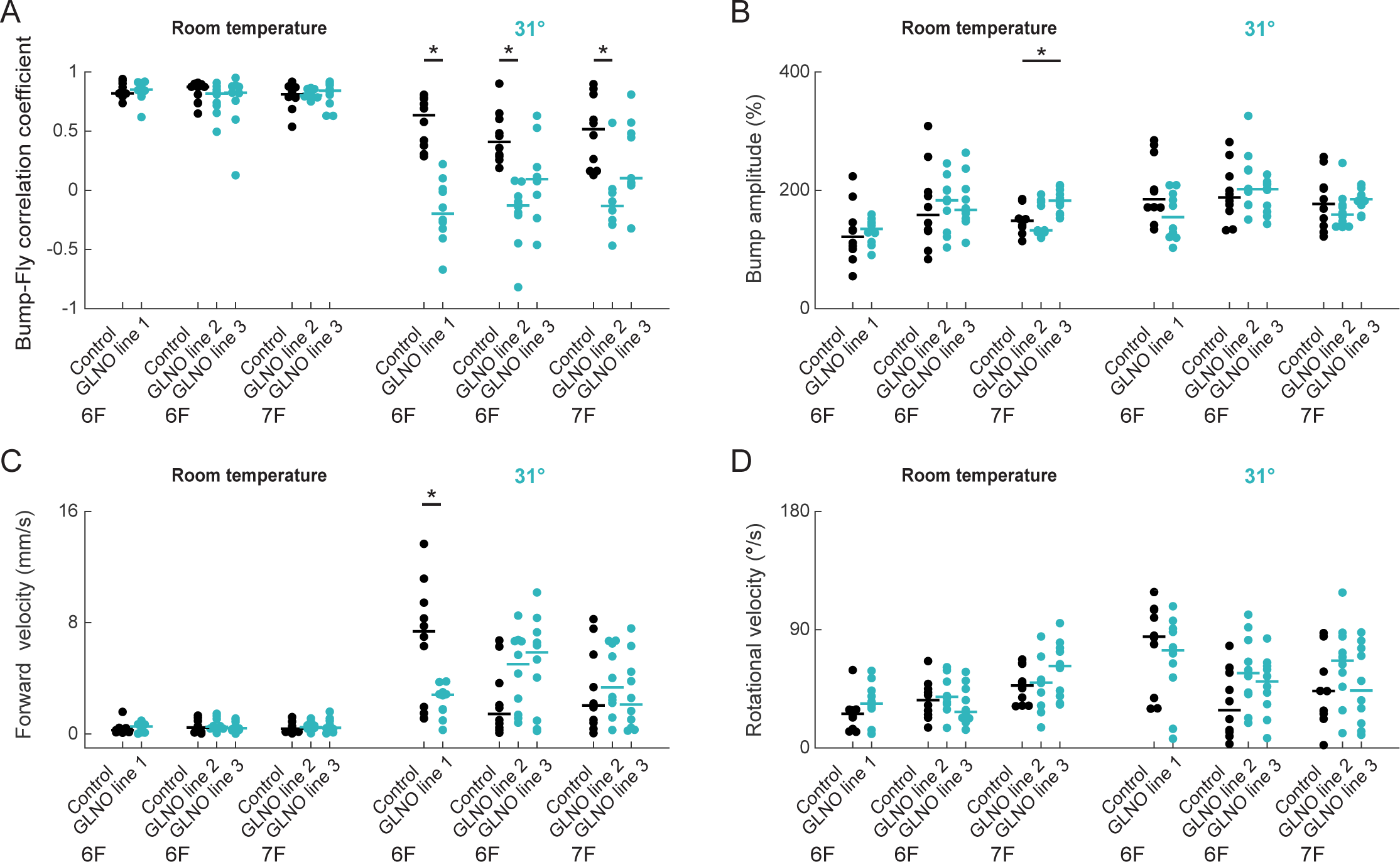
GLNO neurons are required for accurate angular path integration during closed-loop visual experience. **A)** Same as in Figure 2D but for trials where the fly was given closed-loop control of a ∼20° wide bar, which improves the correlation between the fly’s HD and the bump’s position at room temperature (left dots are all closed to 1) but cannot rescue the reduced bump-fly correlation at higher temperatures in shibirets-expressing flies. **B)** Same as in Figure 2D but showing the average bump amplitude. Reducing transmitter release from GLNO neurons does not impact the shape of the bump, consistent with preserved attractor dynamics. **C)** Same as in Figure 2D but showing the flies’ average forward velocity. Higher temperatures increase the flies’ forward velocity but do so equally for shibire^ts^-expressing and control flies. **D)** Same as in Figure 2D but showing flies’ average rotational velocity.

**Figure 3 S1:**
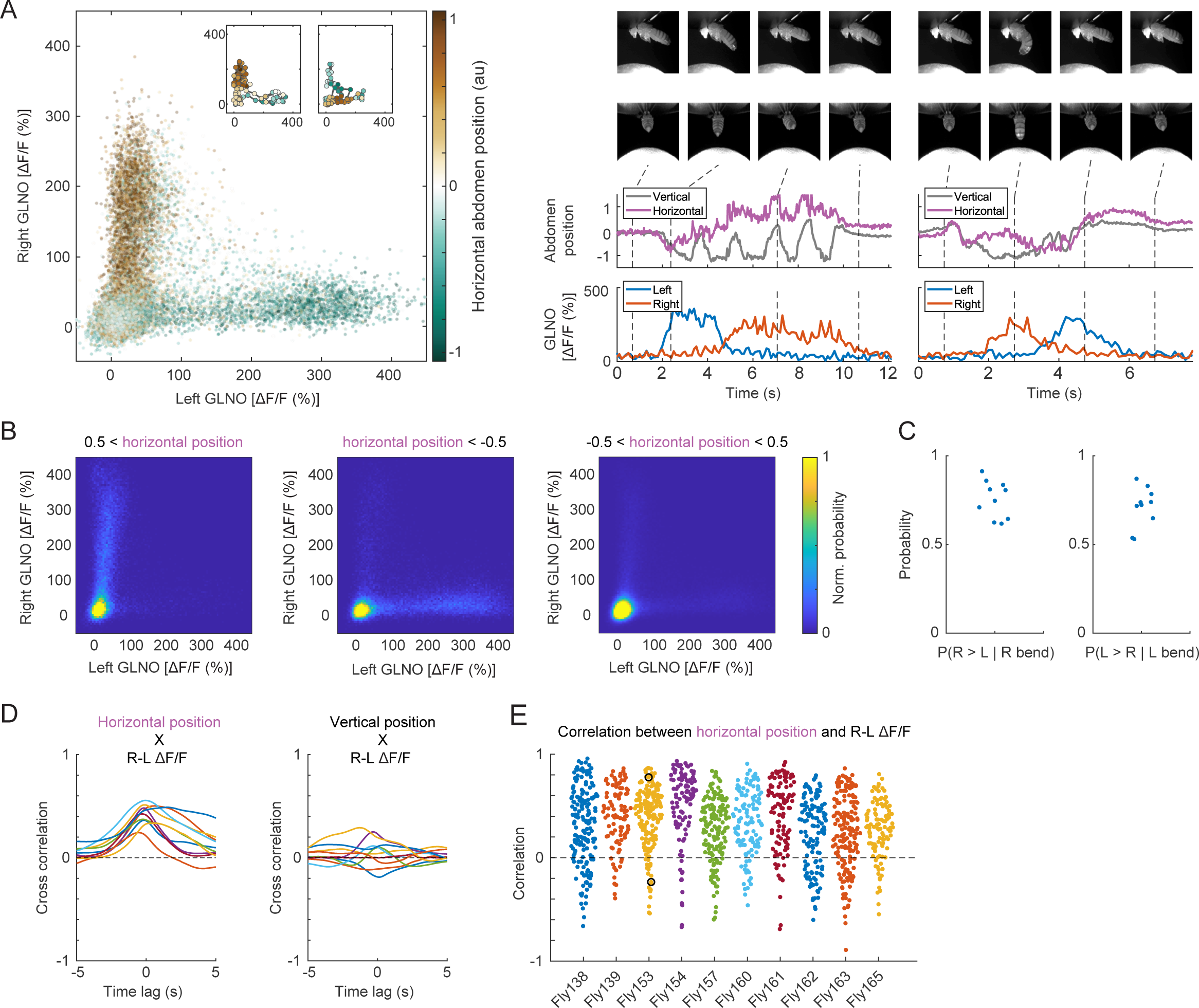
Partial dissociation between GLNO activity and the fly’s proprioceptive state. **A)** Example data from a single fly showing correlation between abdomen bends and GLNO activity in darkness. Flies had their legs removed to disrupt naturalistic proprioceptive activity patterns, as might be experienced during walking. Scatter plot shows the left versus right GLNO activity, color-coded according to the normalized horizonal position of the abdomen (positive values indicate rightward abdomen bends). Notice that rightward abdomen bends tend to be associated with activation of the right GLNO neurons and leftward abdomen bends with activation of the left GLNO neurons, as exemplified by the middle panel, but there are periods where the opposite occurs, as exemplified in the right panel. In addition, most periods with abdomen bends were not associated with GLNO activity (points near the origin). Right panel groups depict two periods of GLNO activity with opposing correlation profiles to the abdomen position (corresponding to the insets in the left panel). Top two rows illustrate the side and rear view of the fly, respectively, at different time points (dashed lines in the lower rows). Bottom two rows depict the abdomen position and GLNO activity changes during the selected periods, respectively. **B)** 2D histograms showing the normalized joint probability of the left and right GLNO activity during periods with large leftward abdomen bends (above 1 std, left panel), large rightward abdomen bends (middle panel), and small amplitude abdomen bends in either direction (right panel). Notice that most abdomen bends do not lead to appreciable activity in the GLNO neurons, suggesting a partial dissociation between GLNO activity and fly’s proprioceptive state. **C)** Beeswarm plot showing the probability that the right GLNO neurons were more active than the left GLNO neurons during a rightward abdomen bend (left panel), and correspondingly for leftward abdomen bends (right panel). **D)** Cross correlations between GLNO activity difference (right – left) and horizontal (left panel) and vertical (right panel) abdomen bends. **E)** Beeswarm plot showing the correlation between GLNO activity difference and horizontal abdomen bends, where each point represents the correlation from a discrete GLNO activity bout from a single fly. Notice that while most bouts are associated with positive correlations, the strength (and sign) of this correlation shows considerable variability. The two black circles for fly 153 correspond to the example data shown in (A).

**Figure 3 S2:**
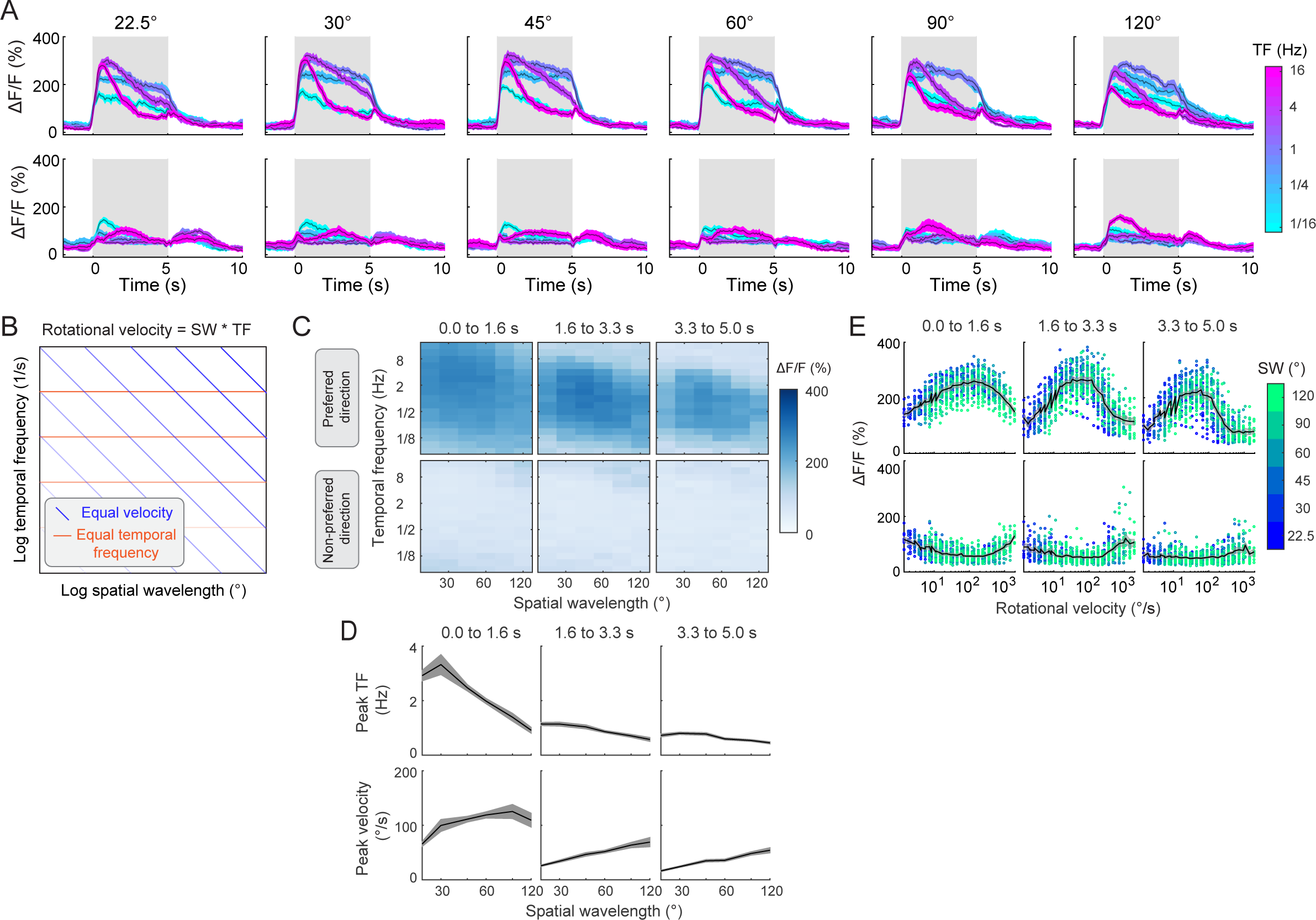
Spatiotemporal tuning profiles of GLNO optic flow responses. **A)** Fly-averaged GLNO responses (+/- s.e.m.) to motion at five different temporal frequencies (marked on heatmap’s y-axis) using square wave gratings with six different spatial wavelength (indicated above each plot) for motion in the preferred (top panels) and non-preferred directions (bottom panels). **B)** Schematic showing two competing models that could describe GLNO optic flow tuning: equal velocity tuning (blue lines) or equal temporal frequency tuning (orange lines; (Creamer et al., 2018)). A stimulus’s rotational velocity is equal to the product of its spatial wavelength (in degrees) and its temporal frequency (in Hz), giving rise to diagonal lines marking stimuli with the same rotational velocity on a log-log plot. Darker blue lines indicate higher rotational velocity and darker orange lines indicate higher temporal frequencies. **C)** 2D heatmap showing fly-averaged GLNO responses to motion stimuli as a function of the stimulus’s spatial wavelength (x-axis) and its temporal frequency (y-axis) for different time windows after stimulus onset (indicated above each plot) and for motion in the preferred (top tow) and non-preferred directions (bottom row). **D)** Fly-averaged GLNO responses (+/- s.e.m.) showing the peak temporal frequency (top panels) and peak rotational velocity (bottom panels) as a function of the stimulus’s spatial wavelength (x-axis). For each column in the preferred direction matrices from (C), we used a Gaussian fit to determine which temporal frequency yields the largest GLNO activation, plotted in the top panels for the same three time windows as in(C). The peak velocity, shown in the bottom panels, was obtained by multiplying the peak temporal frequency by stimulus’s spatial wavelength. **E)** Scatter plot showing the per-trial GLNO response amplitude across all flies, as a function of stimulus’s rotational velocity (x-axis) and spatial wavelength (indicated by the color of the circles) for motion in the neurons’ preferred (top row) and non-preferred (bottom row) directions. The average response (+/- s.e.m.) across flies is shown in black/gray.

